# Inhibition of mucin-type *O-*glycosylation impairs melanogenesis, melanoma growth, and metastatic capacity

**DOI:** 10.64898/2026.02.25.707981

**Authors:** Tanya Jain, Abhiraj Rajasekharan, Simran Kumari Gorai, Aishwarya Rajan, Nagarajan Perumal, Santiswarup Singha, Srinivasa-Gopalan Sampathkumar

## Abstract

Treatment of malignant and metastatic melanoma remains a clinical challenge, despite the success of immune checkpoint blockade and kinase inhibitors, highlighting the need to target complementary pathways. Cancerous cells display a dense glycocalyx, containing tumor-associated carbohydrate antigens, which serves as a shield against immune recognition, confers a survival advantage to the tumor, and reduces intracellular drug entry. On the one hand, interactions between sialoglycans expressed on tumor cell surfaces and sialic acid-binding immunoglobulin-type lectins (Siglecs) on immune cells facilitate immune suppression. On the other hand, hyperactive melanosome and higher melanin content correlate with a poor prognosis in melanoma. The *O*-glycosylation of PMEL17/gp100, overexpressed in melanoma, is indispensable for melanosome architecture, melanin biosynthesis, and melanoma aggressiveness. We show that pharmacological inhibition of O-glycosylation in B16F10 (mouse melanoma) cells using peracetyl *N*-thioglycolyl-D-galactosamine (Ac5GalNTGc, **1a**) decreased both PMEL/17gp100 glycosylation and melanin content, and diminished the Siglec-E engagement to disrupt the sialoglycan-siglec immune-checkpoint through hyposialylation of the cell surface. In C57BL/6J mice, administration of **1a**, but not controls, significantly delayed B16F10 tumor growth, prolonged the survival of mice, and significantly reduced the lung metastases. These findings identify *O*-glycosylation as a dual-mode therapeutic vulnerability, combining melanosome dysfunction and abrogation of siglec-mediated immune evasion in melanoma.

**Statement of Significance:** Sialoglycans on melanoma cells bind Siglecs on immune cells, triggering immunosuppression. Inhibiting *O*-glycosylation with peracetyl *N*-thioglycolyl-D-galactosamine (Ac5GalNTGc, **1a**) caused PMEL17/gp100 hypoglycosylation and abrogated Siglec-E binding, underscoring glycosylation pathways as potential anti-cancer targets.

## Introduction

Specific molecular processes, regulated spatially and temporally, define multicellular development and homeostasis. However, the probability of disrupted homeostasis, dysregulated differentiation, immune exhaustion, epithelial-mesenchymal transition (EMT), and malignant transformation increases exponentially with age, necessitating effective interventions (1). Cancer exemplifies how dysregulated developmental and immune processes drive lethal disease.

Melanoma is an aggressive cancer, characterized by high metastatic potential and poor prognosis. Despite advances in checkpoint immunotherapy (PD-1/PD-L1, CTLA-4/B7) and kinase inhibitors (BRAF V600E), resistance, relapse, and metastasis remain significant challenges (2). Evidence suggests that melanogenesis and melanin accumulation contribute to melanoma aggressiveness, with increased melanin content positively correlated with disease severity and metastatic potential (3).

Melanogenesis occurs within melanosomes, specialized organelles in melanocytes. Melanosomes contain PMEL17/gp100 (premelanosomal protein) and MART-1/Melan-A within their bilayer membrane and luminal compartment, where eumelanin and pheomelanin are synthesized through tyrosinase-driven oxidation of tyrosine and DOPA. These reactions involve reactive quinone intermediates and free radicals, necessitating tight spatial confinement to prevent cytotoxic damage. PMEL17/gp100 forms organized amyloid-like fibrils that promote efficient melanin polymerization and controlled deposition (4).

PMEL17/gp100 is synthesized as a glycosylated precursor and undergoes proteolytic cleavages prior to fibril self-assembly. Both the polycystic kidney disease (PKD) and repeat (RPT) domains of PMEL17/gp100 are critical for maintaining fibrillar melanosomal architecture (5). Its mucin-like domain, enriched in repetitive sequences and dense *O*-glycosylation, likely promotes controlled amyloidogenesis. Melanoma frequently overexpresses melanocyte-specific antigens, including PMEL17/gp100, tyrosinase, and tyrosinase-related proteins 1 and 2 (TYRP1/2) (6). Although most melanosomal proteins are glycoproteins, few studies have explored glycosylation as a therapeutic target for melanoma or pigmentation disorders.

Aberrant glycosylation is a hallmark of cancer, contributing to tumor growth, invasion, and immune evasion. Tumor cells exhibit altered mucin-type *O-*glycosylation (MTOG) patterns, including truncated *O-*glycans like sialyl-Tn-antigen (GalNAcα-Ser/Thr), altered core structures, and increased terminal sialylation (7). Many are recognized as tumor-associated carbohydrate antigens (TACAs). These alterations promote tumor survival by enhancing cell adhesion, resisting apoptosis, masking tumor epitopes from cytotoxic T lymphocytes (CTLs) through altered antigen processing, and engaging immune-inhibitory receptors (8).

Recently, the Siglec-sialoglycan axis emerged as a critical immune checkpoint. Siglecs, expressed on immune cells, bind sialylated glycans on tumor cells and transduce immunosuppressive signals through immunoreceptor tyrosine-based inhibitory motifs (ITIM), dampening anti-tumor responses. Beyond anti-PD-1/CTLA-4, targeted removal of sialic acids using antibody-enzyme conjugates (AEC) has shown promise (9). Büll and coworkers reported that halogen-substituted *N-*acetyl-D-neuraminic acid (NeuAc) derivatives retarded melanoma growth (10). Esko and coworkers reported *O-*glycosylation inhibitors using decoy mechanisms (11). However, both approaches target MTOG elaboration, leaving the initiation steps of *O*-glycosylation largely unexplored.

*N*-acetyl-D-hexosamine (HexNAc) analogues manipulate glycoconjugates through salvage pathways. Analogues bearing keto-, azido-, and alkyne-functionalities enable metabolic glycan engineering (MGE), while halogen-substituted analogues inhibit glycosylation without metabolic incorporation (10,12).

We previously demonstrated that peracetyl *N*-thioglycolyl-D-galactosamine (Ac_5_GalNTGc, **1a**) efficiently inhibits MTOG *in vitro* and *in vivo* (13–15). **1a** is metabolically incorporated into glycoproteins and inhibits both the glycan site occupancy of the Tn antigen (α-GalNAc-Ser/Thr), the first step in MTOG biosynthesis, and subsequent elaboration, inducing hyposialylation. We hypothesized that *O*-glycosylation inhibition would disrupt PMEL17/gp100-dependent melanosome integrity and diminish tumor sialoglycans, potentially impairing Siglec-mediated immune suppression and enhancing anti-tumor immunity (**Fig. 1 A**).

**Figure 1.**
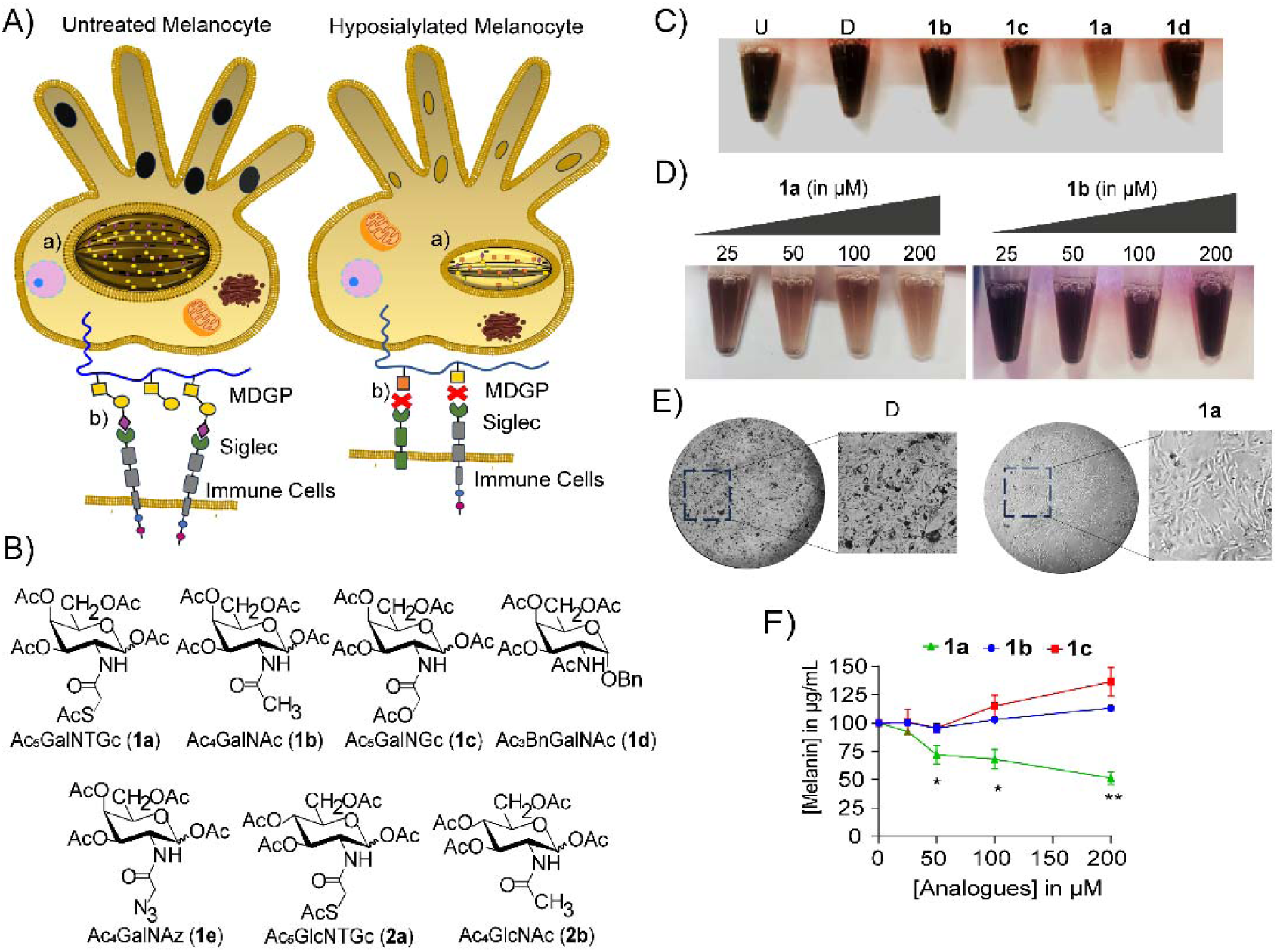
Metabolic glycan engineering using modified GalNAc analogues disrupts melanin biosynthesis. (A) Schematic model illustrating the role of *O*-glycosylation in melanosome structure and melanogenesis. Under physiological conditions, melanosomes contain organized amyloid-like fibrils of PMEL/gp100 that provide a scaffold for melanin deposition and confer ellipsoid, sheet-like architecture. (a) Inhibition of *O*-glycosylation by **1a** disrupts melanosome integrity and melanogenesis, leading to defective melanosomes and decreased pigmentation. In cancer cells, hypersialylation promotes immune evasion via Siglec engagement; **1a**-induced hyposialylation may attenuate this Siglec-sialic acid axis. (B) Chemical structures of GalNAc and GlcNAc analogues used in this study: Ac_5_GalNTGc (**1a**), featuring a thioglycolyl group in the *N-*acyl moiety; Ac_4_GalNAc (**1b**), the peracetylated form of the wild-type GalNAc analogue; Ac_5_GalNGc (**1c**), featuring a glycolyl group in the *N*-acyl moiety; Ac3BnGalNAc (**1d**), an *O*-glycan inhibitor distinguished by its benzyl group; Ac4GalNAz (**1e**), featuring an azide moiety; Ac_4_GlcNAc (**2b**), the peracetylated form of the wild-type GlcNAc analogue; and Ac_5_GlcNTGc (**2a**), the C4 epimer of **1a**. (C) Visual pigmentation of B16F10 cells suspended in 1× PBS after treatment with GalNAc analogues (100 µM, 48 h). (D) Dose-dependent pigmentation analysis of B16F10 cells suspended in 1× PBS treated with **1a** or **1b** (25-200 µM, 48 h). (E) Phase-contrast images (10×) of B16F10 cells treated with D (vehicle) or **1a**, showing intracellular melanin granules. (F) Quantification of melanin content following treatment with increasing concentrations (25, 50, 100, and 200 µM) of GalNAc analogues. Data represent mean ± SD (standard deviation) of three biological replicates. Statistical significance was determined by using one-way analysis of variance (ANOVA). *P <0.05, **P <0.01 vs. D). U, untreated; and D, DMSO (vehicle). Monosaccharide symbols follow the Symbol Nomenclature for Glycans (SNFG)(49).

Herein, we report that **1a** profoundly alters melanoma biology. Treatment decreased PMEL17/gp100 glycosylation, melanin content, and global sialylation, impairing migration and invasion in B16F10 and B16F10-Luc2 mouse melanoma cells. In a syngeneic model, **1a** significantly retarded tumor growth, reduced lung metastasis, and prolonged survival. Collectively, these findings establish *O*-glycosylation as a promising therapeutic target and support carbohydrate-based small-molecule inhibitors for combinatorial adjuvant and neoadjuvant therapy.

## Results

Mucin-type *O*-glycans are abundant in mucins and mucin-domain glycoproteins (MDGP) (e.g., CD43, CD162/PSGL-1), particularly within linker regions of membrane and secreted glycoproteins (e.g., IgM, IgA). MTOG synthesis is initiated in the Golgi by polypeptide GalNAc transferases (ppGalNAcTs), utilizing UDP-GalNAc to form the Tn-antigen (16) (**Fig. S1 A**). The Tn-antigen is further elaborated into the Thomson-Friedenreich antigen (TF- or T-antigen), sialyl-Tn, and disialyl-T structures by galactosyl- and sialyl-transferases.

To investigate structure-activity relationships, we employed a panel of GalNAc analogues, *viz*., peracetyl *N-*thioglycolyl-D-galactosamine (Ac_5_GalNTGc, **1a**; thiol-containing) (17), peracetyl *N*-acetyl-D-galactosamine (Ac_4_GalNAc, **1b**; wild-type control) (13), peracetyl *N*-glycolyl-D-galactosamine (Ac_5_GalNGc, **1c**; *N-*glycolyl analogue) (13), peracetyl benzyl-*N*-acetyl-D-galactopyranoside (Ac_3_BnGalNAc, **1d**; substrate decoy inhibitor) (13), peracetyl *N*-azidoacetyl-D-galactosamine (Ac_4_GalNAz, **1e**; reporter analogue) (18), peracetyl *N*-thioglycolyl-D-glucosamine (Ac_5_GlcNTGc, **2a**; C-4 epimer of **1a**) (15), and peracetyl *N*-acetyl-D-glucosamine (Ac_4_GlcNAc, **2b**; C-4 epimer of **1b**) (19) (**Fig. 1 B**).

### Inhibition of *O-*glycosylation decreases melanin biosynthesis in B16F10 cells

To study *O-*glycosylation in melanogenesis, we employed B16F10 mouse melanoma cells, a widely used *in vitro* and *in vivo* model (20). Following treatment with GalNAc analogues: **1a**, **1b**, **1c**, or **1d** (100 µM, 48 h), harvested intact cell suspensions revealed that **1a**-treated cells showed no visible pigmentation compared to other groups (**Fig. 1 C**). This depigmentation was specifically induced by **1a**, but not by **1b**, in a dose-dependent manner (25-200 µM) (**Fig. 1 D**). Phase contrast microscopy confirmed abundant visible melanin granules in DMSO (vehicle) controls, which were notably absent in **1a**-treated cells (**Fig. 1 E**). Quantitative spectrophotometric analysis at 405 nm confirmed a ∼ 75 % decrease in melanin content following **1a** treatment, while DMSO, **1b**, **1c**, and **1d** resulted in only modest decreases of 10 %, 25 %, 10 % and 35 %, respectively, compared to the untreated control (**Fig. S1, B and C**). This inhibition was dose-dependent (0-200 µM) for **1a**, whereas higher concentrations of **1b** and **1c** slightly increased melanin content (**Fig. 1 F**).

To evaluate potential cytotoxicity, MTT assays showed that **1a** and **1b** maintained 90 % cell viability up to 100 µM, whereas **1c** caused moderate toxicity (∼ 40 % decrease in viability) at 100 µM (**Fig. S1 D**). Trypan blue exclusion and Coulter counting confirmed comparable viable cell numbers for DMSO, **1b**, **1d**, and **1a**-treated cultures, with a ∼ 20 % decrease observed only for **1c** (**Fig. S1, E and F**). Carboxyfluorescein diacetate succinimidyl ester (CFSE) dilution analysis by flow cytometry revealed comparable proliferation rates across all treatment groups at 48 h, indicating no cell cycle arrest in **1a**-treated cells (**Fig. S1 G**). Together, these results demonstrate that **1a** potently inhibits melanin biosynthesis independently of cytotoxic or antiproliferative effects, distinguishing it from controls and other analogues.

### Disruption of PMEL17/gp100 glycosylation by 1a

Because **1a** induces hyposialylation of the *O-*glycosylated protein CD43, we hypothesized it would similarly perturb PMEL17/gp100, a melanosomal structural protein (13). Mouse PMEL17/gp100/silv is a 626-amino acid multi-domain protein consisting of a signal peptide (SP), *N*-terminal region (NTR), polycystic kidney disease (PKD) domain, repeat (RPT) domain, cleavage site (CS), kringle-like domain (KLD), and *C*-terminal domain (CTD) (**Fig. 2 A**).

**Figure 2.**
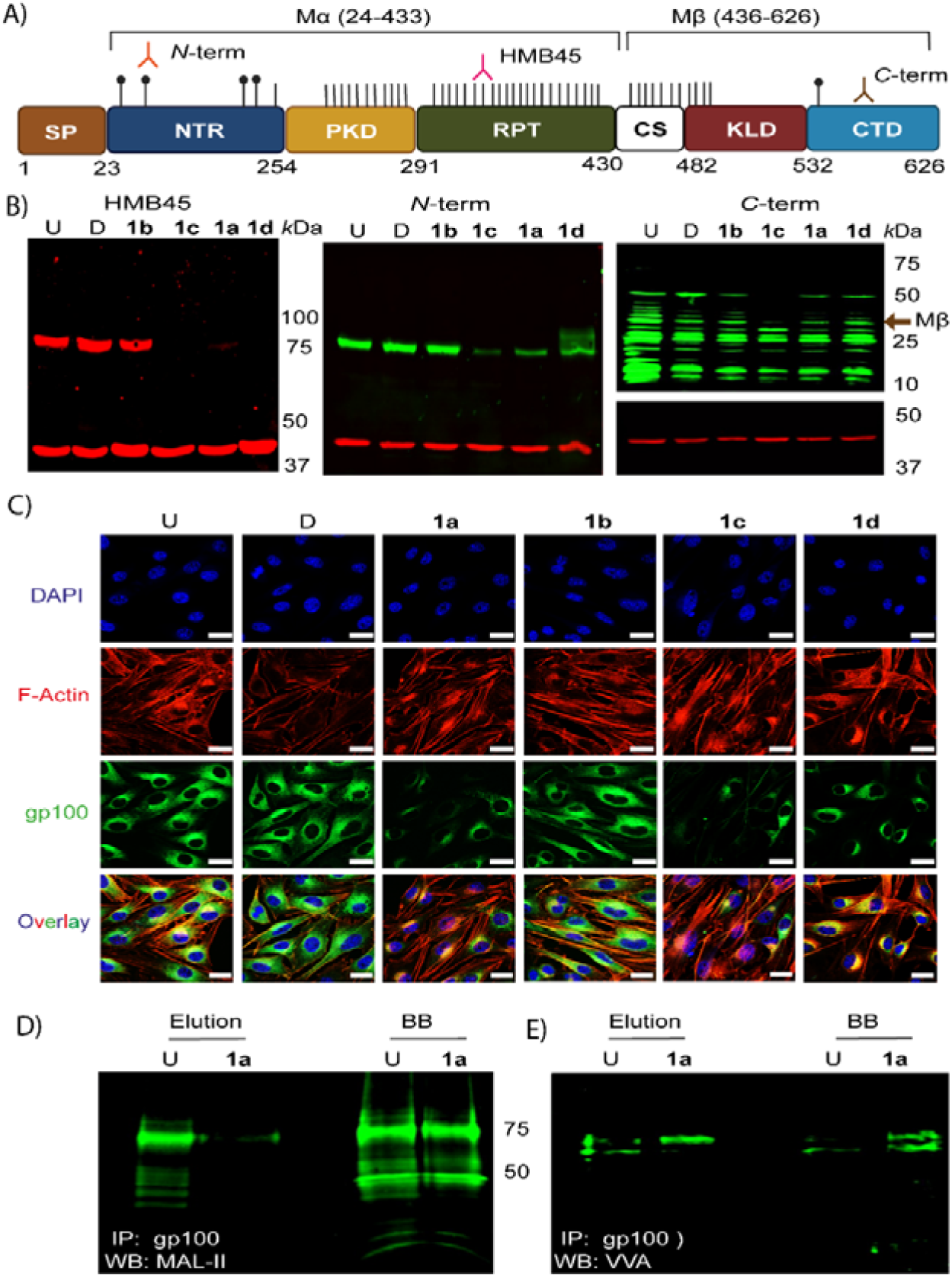
Modulation of PMEL/gp100 glycoforms upon analogue treatment in B16F10 cells. (A) Schematic representation of predicted glycosylation sites on mouse PMEL/gp100. The 626-amino acid protein is modified by four *N*-linked glycans and ∼60 potential *O*-linked glycans, with 56 sites clustered within the PKD and RPT domains that contribute to fibril formation. SD, signal domain; NTD, *N*-terminal domain; PKD, polycystic kidney disease domain; RPT, repeat domain; CS, cleavage site; KRG, kringle-like domain; TM, transmembrane; and CTD, *C*-terminal domain. Amino acid positions marking the start of each domain are indicated below. *O*-glycosylation sites are indicated by black lines and *N*-glycosylation sites by lines with circles. Binding regions for glycan-dependent (HMB45), *N*-terminal, and glycan-independent *(C*-terminal) PMEL/gp100 antibodies are indicated. (B) Duplex immunoblot analysis of PMEL/gp100 in B16F10 cells treated with GalNAc analogues (100 µM, 48 h). Blots were probed with glycan-dependent HMB45 (left panel) (upper red band), N-term (middle panel) (upper green band), or glycan-independent C-term (right panel) (upper green band), with β-actin as a loading control (lower red band). Images were acquired on an Amersham Typhoon fluorescent imager. Blots shown are representative of three biological replicates (BR). (C) Confocal imaging of B16F10 cells treated with GalNAc analogues (100 µM, 48 h) and stained for PMEL/gp100 (*N*-term, green), F-actin (phalloidin, red), and nuclei (DAPI, blue). Imaging was performed using 63× oil immersion on a Zeiss LSM 980 confocal microscope. The images shown are representative of three BRs, with two technical replicates per BR. Each replicate was conducted on a single cover slip, with five fields imaged per coverslip. Scale bar: 5 µm. (D-E) Lectin blot analysis of immunoprecipitated PMEL/gp100 eluted by 0.2 M Glycine (pH 2.6) showing (D) reduced MAL II binding and (E) increased VVA binding upon **1a** treatment, indicating loss of terminal α2,3-linked sialic acids and exposure of the Tn antigen, respectively. Both the eluted fraction and the boiled beads (BB) fractions are shown. The data are representative of three BR. MAL II (*Maackia amurensis* Lectin II); VVA (*Vicia villosa* agglutinin); U, untreated; D, DMSO (vehicle); **1a**, Ac_5_GalNTGc; **1b**, Ac_4_GalNAc; **1c**, Ac5GalNGc; and **1d**, Ac3BnGalNAc.

Following synthesis, PMEL17/gp100 undergoes extensive post-translational glycosylation and proteolytic processing, generating multiple functional fragments, including the *N-*terminal Mα, Mα-Mβ, PKD-RPT, and *C-*terminal Mβ fragments (21). The PKD-RPT domain is particularly critical for the formation of the organized, amyloid fibrillar architecture of melanosomes. Several gp100-derived peptides presented on MHC molecules have been explored as melanoma vaccine candidates, including gp100_25-33_ (KVPRNQDWL) and gp100_44-59_ (WNRQLYPEWTEAQRLD), which originate from NTR, as well as gp100_209-217_ (ITDQVPFSV) and gp100_280-288_ (YLEPGPVTA), derived from the PKD domain; predominantly arise from non-glycosylated regions, suggesting dense glycosylation may mask potential antigens (22,23).

*In silico* analysis (Uniprot Q60696) using NetNGlyc and NetOGlyc predicted four *N*-glycan sites and 60 *O*-glycan sites, mostly clustered within the PKD and RPT domains (24,25) (**Fig. 2 A**). The dense clustering of *O-*glycans is proposed to create a bottle-brush-like structure essential for organized fibril formation and its stabilization; the RPT domain specifically carries core 1 *O*-glycans with Tn, sialyl-Tn, sialyl-T, and disialyl-T structures (26). To assess glycosylation changes, B16F10 cell lysates treated with GalNAc analogues (100 µM, 48 h) were analyzed by western blotting using three antibodies: (i) anti-gp100 (*N-*term); (ii) anti-gp100 (HMB45), binding RPT-domain sialoglycans, and (iii) anti-gp100 (*C-*term) (26).

Strikingly, anti-gp100 (HMB45) epitopes were abrogated upon treatment with **1a**, **1c**, and **1d**, whereas DMSO (vehicle) and **1b** showed no change compared to the untreated control (**Fig. 2 B**, left panel). Anti-gp100 (*N-*term) blots showed a significant decrease in the 75 *kDa* bands upon **1a** and **1c** treatment, while **1d** treatment produced diffuse higher molecular-weight bands above 75 *kDa*, and DMSO and **1b** treatment resulted in no changes (**Fig. 2 B**, middle panel). Although **1c** altered glycoforms, it also induced cytotoxicity without decreasing melanin content. In the anti-gp100 (*C-*term) blot, **1a**, **1b**, **1d**, and DMSO showed no change in the 28 *k*Da band intensity. Importantly, **1a** did not alter *C-*terminal protein levels, indicating its effect is restricted to post-translational glycosylation rather than expression or processing (**Fig. 2 B**, right panel). In contrast, **1c** treatment resulted in a clear decrease in the 28 *k*Da band, indicating decreased PMEL17/gp100 protein stability or enhanced degradation under these conditions.

The effects of **1a** were dose-dependent and reversible. Suppression of glycosylation became evident at 50 µM and was maximal at 100-200 µM, as detected by both anti-gp100 (*N*-term and HMB45) antibodies (**Fig. S2, B and C**). Time-course analysis (100 µM, 0-96 h) showed maximal suppression at 48-72 h, with full recovery by 96 h, reflecting normal biosynthetic turnover of glycoproteins, analogue dilution, and endogenous GalNAc restoration (**Fig. S2 D**). Confocal microscopy confirmed the loss of anti-gp100 (*N-*term and HMB45) epitopes upon treatment with **1a**, **1c**, or **1d**, but not with D or **1b** (**Fig. 2 C, and Fig. S2 A**), while F-actin staining confirmed cellular morphology was unaffected. To show that **1a** directly alters PMEL17/gp100, the protein was immunoprecipitated from B16F10 lysates. Lectin blotting of immunoprecipitated PMEL17/gp100 revealed a marked reduction in MAL-II (*Maackia amurensis* lectin-II) binding upon **1a** treatment, compared to control, indicating a specific loss of terminal α2→3-linked sialic acids on the protein (**Fig. 2 D**). Conversely, VVA (*Vicia villosa* agglutinin) binding increased, indicating exposure of the underlying Tn antigen (GalNAcα-Ser/Thr) (**Fig. 2 E**). Notably, lectin blotting of the boiled bead fraction, which captures all antibody-bound PMEL17/gp100, including strongly bound or incompletely eluted species, exhibited the same trend, confirming robust changes independent of elution efficiency. These results provide direct molecular evidence that **1a** inhibits the *O-*glycan elongation on PMEL17/gp100. Mechanistically, metabolic incorporation of **1a** likely perturbs glycopeptide-selective core 1 β1→3-galactosyltransferase (C1GalT1) activity, impairing the coordinated elongation of adjacent *O-*glycosylation sites, decreasing glycan density, and causing protein hyposialylation (15).

### Selective disruption of melanosomal glycoprotein enzymes by 1a while sparing structural proteins

To define the selectivity of **1a** for *O-*glycosylated proteins, we analyzed melanosomal proteins with contrasting glycosylation profiles. Tyrosinase is a critical *O-*glycosylated enzyme required for the initiation and regulation of melanin biosynthesis, and its inhibition underlies the mechanism of several anti-melanogenic agents. Tyrosinase possesses five predicted *N-*glycan and four *O-*glycan sites (Uniprot P11344) based on *in silico* NetNGlyc and NetOGlyc analysis (27). Western blotting revealed a pronounced reduction in tyrosinase levels following **1a** treatment, but not upon treatment with DMSO, **1b**, or **1c**, supporting a requirement for intact glycosylation for maintaining tyrosinase stability (**Fig. 3 A**).

**Figure 3.**
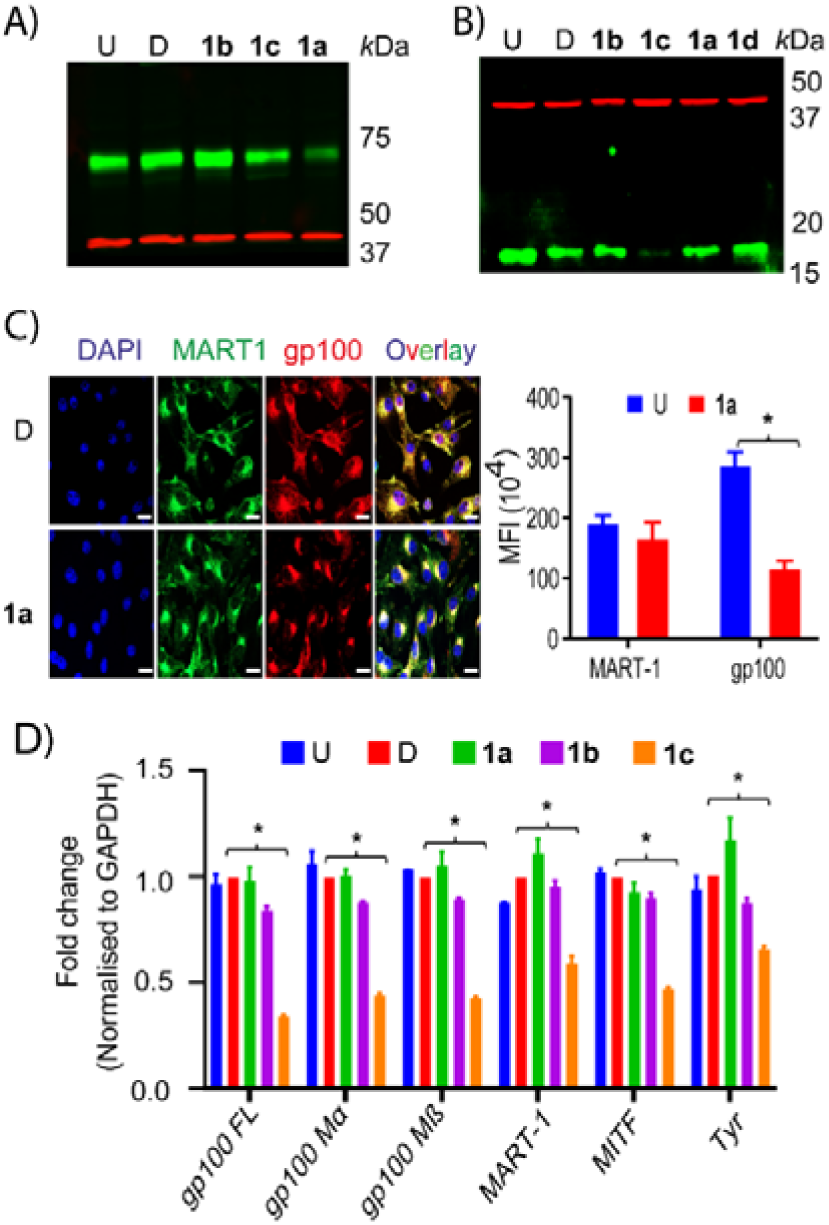
Selective post-translational effects of GalNAc analogues on melanosomal proteins. (A) Effect of GalNAc analogues on tyrosinase levels in B16F10 cells. Cells were treated with GalNAc analogues (100 µM, 48 h), lysed, and probed with anti-tyrosinase antibody (upper green bands) and anti-β-actin (lower red bands; loading control). (B) Effect of GalNAc treatment on MART-1 levels in B16F10 cells. Cells were treated with GalNAc analogues (100 µM, 48 h), lysed, and probed with anti-MART-1 antibody (lower green bands) and anti-β-actin (upper red bands; loading control). (C) Confocal imaging of B16F10 cells either left untreated or treated with **1a** (100 µM, 48 h), fixed, permeabilized, and stained for MART-1 antibody (green), PMEL/gp100 (*N*-term, red), and nuclei (DAPI, blue). Imaging was performed using a Zeiss LSM 980 confocal microscope with a 63× objective. The merged images show colocalization of MART-1 with PMEL/gp100 (yellow). The images shown are representative of three BRs, with two technical replicates per BR. Each replicate was conducted on a single coverslip, with five fields imaged per coverslip. Scale bar: 5 µm. (D) RT-qPCR analysis of expression of melanosomal-specific genes upon GalNAc analogue treatment in B16F10 cells. Transcript levels of PMEL/gp100 (full-length, Mα, and Mβ), MART-1, MITF, and tyrosinase were quantified following analogue treatment (100 µM, 48 h). Error bars represent the standard deviation of three BR with three technical replicates for each BR. Statistical analysis was performed using two-way ANOVA. P values: *<0.05 compared to vehicle, D. U, untreated; D, DMSO (vehicle); **1a**, Ac_5_GalNTGc; **1b**, Ac_4_GalNAc; **1c**, Ac_5_GalNGc; **1d**, Ac_3_BnGalNAc; and FL, Full length.

To validate this specificity, we examined MART-1/Melan-A, a non-glycosylated melanosomal structural protein that facilitates melanosome maturation and stabilizes PMEL17/gp100 (28). In contrast to PMEL/gp100, which harbors ∼60 predicted *O-*glycan sites and four *N-*glycan sites, MART-1 is predicted to carry neither *O-* nor *N-*glycans. Western blotting and confocal microscopy with an anti-MART-1 antibody showed that MART-1 expression (∼16 kDa) remained unaffected by **1a** (100 µM, 48 h) and by other non-toxic controls (**Fig. 3, B** and **C**). This indicates that **1a** does not affect non-glycosylated protein levels. In contrast, **1c** treatment declined MART-1 expression, consistent with its cellular toxicity rather than selective *O-*glycosylation inhibition.

To exclude transcriptional or splicing alterations, transcript levels were measured by qRT-PCR for full-length PMEL17/gp100 (FL), Mα, and Mβ fragments. Across all primer sets, the **1a** treatment did not significantly change PMEL17/gp100 transcript levels compared to untreated and DMSO controls (**Fig. 3 D**). Similarly, transcripts of other key melanogenic regulators, including MART-1, MITF, and tyrosinase, remained unchanged upon **1a** or **1b** treatment, whereas **1c** exhibited a modest decrease across several transcripts, consistent with its partial cytotoxic effects (**Fig. 3 D**). These data demonstrate that **1a** does not alter the transcription of melanosomal structural or regulatory genes.

Collectively, **1a** acts selectively at the post-translational level by perturbing the glycosylation of tyrosinase and PMEL17/gp100 while sparing non-glycosylated proteins like MART-1. This selectivity confirms the direct mechanistic link between *O-*glycosylation inhibition and melanogenesis blockade.

### 1a induces hyposialylation, disrupts Siglec-E recognition, and arrests the migration and invasion of melanoma cells

Melanin levels are tightly linked to melanoma biology, where altered pigmentation states correlate with tumor progression, immune evasion, and metastatic potential (29). We examined whether **1a**-induced changes extended to global cellular glycosylation using a lectin panel: MAL-II (NeuAcα2→3Gal), *Sambucus nigra* agglutinin (SNA) (NeuAcα2→6Gal/GalNAc), wheat germ agglutinin (WGA) (α2→3 and α2→6 sialylated residues), and VVA (Tn-antigen, GalNAcα-Ser/Thr) (**Fig. 4 A**).

**Figure 4.**
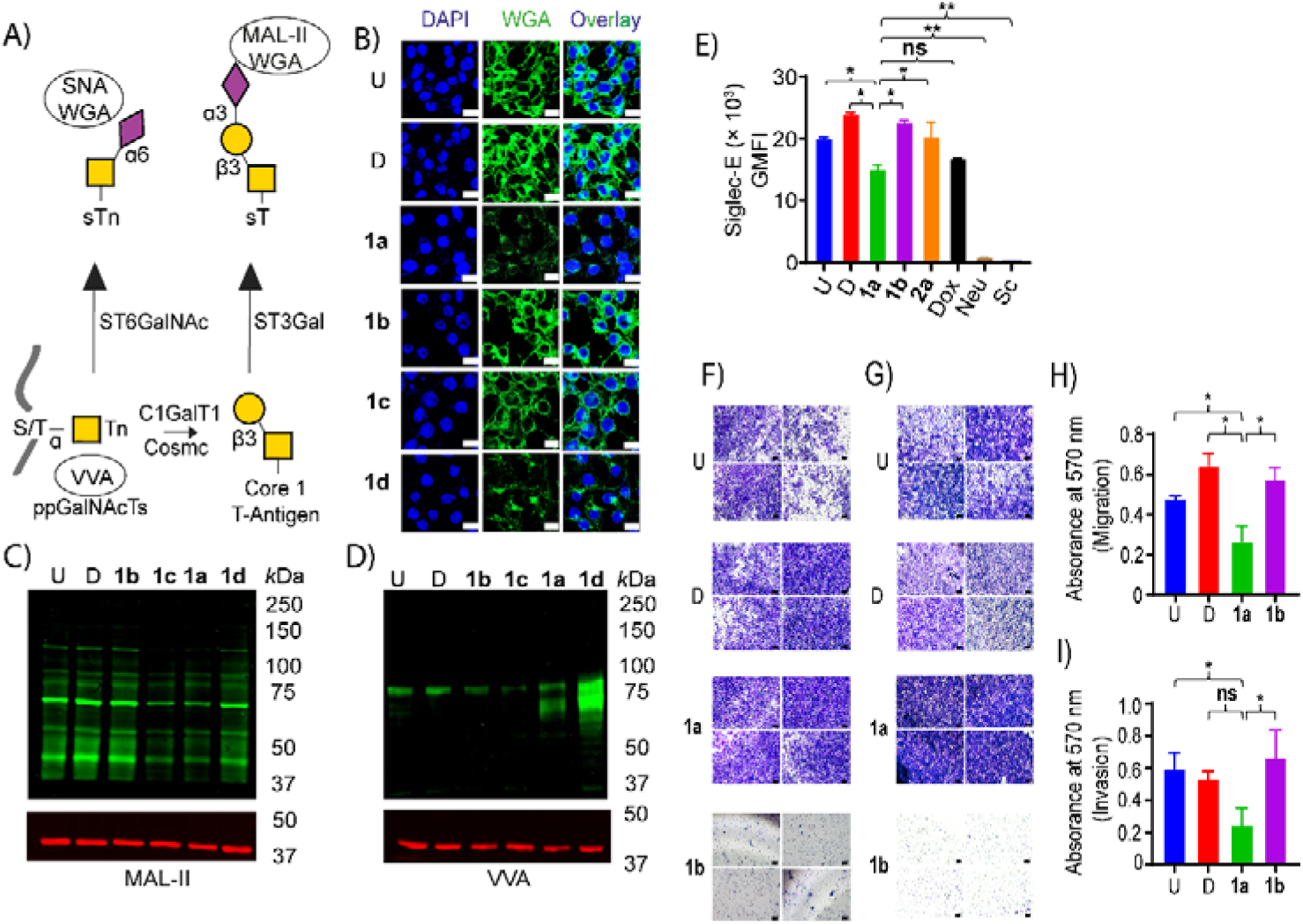
1a causes hyposialylation, decreases Siglec-E binding, and suppresses melanoma migration and invasion. (A) Schematic representation of the binding specificity of lectins used in this study. (B) Confocal microscopy of B16F10 cells left untreated (U) or treated with D, **1b**, **1c**, **1a,** or **1d** (100 µM, 48 h), fixed, permeabilized, and stained with fluorescein-conjugated wheat germ agglutinin (WGA) (green) and DAPI (blue). Imaging was performed using a Zeiss LSM 980 confocal microscope with a 63× objective. The images shown are representative of three BRs, with two technical replicates per BR. Each replicate was performed on one cover slip, with five fields imaged per cover slip. Scale bar: 5 µm. (C-D) Lectin blot analysis of B16F10 cells treated with GalNAc analogues (100 µM, 48 h), lysed and probed with lectins: (C) *Maackia amurensis* lectin-II (MAL-II) (α2→3 sialic acids), and (D) *Vicia villosa* agglutinin (VVA) (Tn-antigen). β-actin blots are shown as loading controls. Blots are representative of three BR. (E) Flow cytometric analysis of Siglec-E binding to B16F10-Luc2 melanoma cells (1.0 × 10^6^) left untreated (U) or treated with vehicle (DMSO), **1b** (100 µM), **1a** (100 µM), **2a** (100 µM), or Dox (100 nM) for 48 h. Cells treated with Neuraminidase from *Vibrio cholerae* (20 milliunits/mL) for 1 h were used as additional controls. Cells were gated sequentially to exclude debris and doublets (**Supporting** Figure 3G). Siglec-E-Fc binding was quantified as geometric mean fluorescence (GMFI) of the Alexa Fluor 488(AF488)-positive cells. Error bars indicate the SD of two BR with two technical replicates each. Statistical significance was determined by one-way ANOVA with Tukey’s multiple comparisons test. P-value * ≤ 0.05, and P-value ** ≤ 0.0001. (F-I) Transwell migration and invasion assays of B16F10-Luc2 (1.0 10^5^) cells left untreated (U) or treated with vehicle (DMSO), **1b** (100 µM), or **1a** (100 µM). Representative images of cells (F) migrated and (G) invaded to the basal side of the filters, fixed, and stained with 0.25 % crystal violet, are shown. [Scale bar: 100 µm, images shown are representative of four fields from one replicate, 10× objective]. Quantification of migration (H) and invasion (I) was performed by solubilizing the cells that migrated to the basal side in DMSO and measuring absorbance at 570 nm. Error bars represent the SD of three BR. Statistical significance was determined using one-way ANOVA with Tukey’s multiple comparisons test. P-value * ≤ 0.05. U, untreated; D, DMSO (vehicle); **1a**, Ac5GalNTGc; **1b**, Ac4GalNAc; **1c**, Ac5GalNGc; **1d**, Ac3BnGalNAc; **2a**, Ac5GlcNTGc; Dox, Doxorubicin; and **Neu**, neuraminidase. Monosaccharide symbols follow the Symbol Nomenclature for Glycans (SNFG) (49).

Confocal microscopy of B16F10 melanoma cells incubated with GalNAc analogues (100 µM, 48 h) and stained with FITC-conjugated WGA revealed robust cell-surface sialylation on untreated, DMSO, and **1b**-treated controls. In contrast, **1a**-treated cells displayed significantly decreased WGA staining, confirming a decreased cell-surface sialoglycan abundance (**Fig. 4 B**). Treatment with **1c** or **1d** resulted in less pronounced or modest decreases, respectively.

Lectin blotting confirmed these findings. Following analogue treatment (100 µM, 48 h), MAL-II epitopes (α2→3 sialic acids) were markedly decreased in **1a** and **1c** conditions, whereas **1d** showed minimal change relative to untreated, DMSO, and **1b** controls (**Fig. 4 C**). Reciprocally, VVA binding (Tn-antigen) was significantly increased in **1a**-treated cells and was even more pronounced with **1d**, while remaining largely unchanged in DMSO, **1b**, or **1c** conditions (**Fig. 4 D**). Together, these findings indicate that loss of terminal sialylation unmasks underlying Tn-antigen structures, resulting in increased VVA accessibility. Notably, **1d** induced robust Tn accumulation without a decrease in MAL-II, suggesting it acts as a decoy for immature *O-*glycan structures without affecting pre-existing sialoglycans.

To quantify surface-level glycosylation in live cells, we performed flow cytometry. SNA binding did not change upon **1a** treatment compared with controls, confirming intact α2→6-linked sialic acids on the cell surface. Correspondingly, VVA binding increased by 20 % upon **1a** treatment, indicating enhanced exposure of Tn-antigens (**Fig. S3 A, B, and C**). Importantly, neuraminidase treatment did not further increase VVA signals, confirming that **1a** induces genuine accumulation of Tn antigen due to impaired *O-*glycan elongation, rather than secondary exposure of underlying structures following enzymatic removal of sialic acids.

Having established that **1a** treatment induces hyposialylation and increases Tn-antigen exposure, we investigated the functional consequences of decreased sialoglycans on immune recognition. Siglecs constitute a family of immune receptors that recognize sialylated glycans and activate immunosuppressive signals through ITIM (9). Among murine receptors, Siglec-E, expressed predominantly on macrophages and neutrophils, promotes B16F10 melanoma and immune evasion by engaging tumor sialoglycans (9).

To assess whether **1a**-induced hyposialylation functionally disrupts this axis, we measured recombinant Siglec-E-Fc binding to B16F10-Luc2 cells by flow cytometry (100 µM, 48 h). Robust Siglec-E-Fc binding occurred in untreated cells, consistent with high basal surface sialoglycan expression. Treatment with vehicle (D) and **1b** (wild-type control) resulted in a modest increase of 15 %, which likely reflects non-specific effects of DMSO exposure and mild perturbations in cellular metabolism due to the metabolic processing of acetylated monosaccharides on overall glycan biosynthesis (**Fig. 4 E, and Fig. S3 D**). Strikingly, **1a**-treatment decreased Siglec-E-Fc binding by 35 % compared to vehicle, demonstrating potent disruption of the Siglec-sialoglycan checkpoint.

In contrast, **2a** produced a modest 10 % increase in Siglec-E-Fc binding, suggesting that non-mucin sialoglycans may not contribute to Siglec-E engagement. Doxorubicin (Dox), a cytotoxic control, resulted in a 25 % decrease, likely reflecting non-specific alterations in surface glycoproteins under cellular stress rather than selective glycosylation inhibition. Enzymatic desialylation with neuraminidase yielded a 95 % decrease, validating assay specificity and confirming the central role of sialic acids in Siglec-E recognition. These results establish that **1a** treatment functionally disrupts the Siglec-sialoglycan immune checkpoint by decreasing tumor cell surface sialylation.

Finally, we evaluated the effect of **1a** on melanoma migration and invasion using Boyden chamber assays. Cells treated with the vehicle (D) or **1b** (wild-type control) showed robust migratory capacity comparable to the untreated control (**Fig. 4, F** and **H**). In striking contrast, **1a**-treatment resulted in a drastic 50 % decrease in migratory cells at 100 µM, with a similar inhibition evident at 50 and 200 µM concentrations (**Fig. S3 E**). Similarly, in a Matrigel-coated invasion assay, vehicle and **1b**-treated cells showed invasive behavior similar to that of the untreated control (**Fig. 4, G** and **I**). However, **1a**-treatment resulted in a striking 60 % decrease in invasive capacity at 100 µM, with comparable inhibition observed at both 50 and 200 µM concentrations (**Fig. S3 E**). These findings demonstrate that **1a**-mediated *O-*glycosylation inhibition significantly impairs the migratory and invasive potential of melanoma cells.

### Selective anti-metastatic transcriptional reprogramming by 1a with minimal off-target effects

To investigate the molecular basis of the observed anti-invasive effects, we first employed a targeted RT-qPCR array profiling 84 metastasis-associated genes in B16F10 cells treated with vehicle (D) or **1a** (100 µM, 48 h). Expression of most genes (70 of 84) was unaffected, indicating no broad transcriptional disruption. Notably, **1a** significantly upregulated tumor suppressors Ephb2 (ephrin type-B receptor 2) (2.2-fold) and Itga7 (integrin subunit alpha 7) (2.0-fold), while simultaneously downregulating pro-metastatic regulators, MMP10 (matrix metalloproteinase-10) and Mycl1 (V-myc myelocytomatosis viral oncogene homolog 1) by ∼50 % (0.5-fold) (30) (**Fig. S4, A and B**). This transcriptional signature aligns with the inhibition of migration and invasion, supporting a functional link between *O-*glycosylation inhibition, disrupted Siglec-sialoglycan interactions, and suppression of metastatic phenotypes *in vitro*.

To evaluate global specificity and potential off-target effects of **1a**, we performed unbiased RNA-seq analysis on vehicle- and **1a**-treated cells. Applying strict filters (P_adj_ <0.05, |log_2_Fc|>1.5), only 35 genes out of 16,359 protein-coding genes were differentially expressed (15 upregulated, 20 downregulated) (**Fig. S4, C-J**). Crucially, canonical melanocyte genes, including PMEL17/gp100, tyrosinase, TYRP1, or MITF, did not meet differential expression thresholds, confirming that **1a** does not perturb the core melanocytic transcriptional program.

Differentially expressed genes identified by RNA-seq showed modest enrichment for thiol-mediated redox homeostasis and glutathione (GSH/GSSG) metabolism pathways (e.g., Hmox1, Nqo1, Gsta3, and Slc7a11), suggesting a mild adaptive response involving glutathione buffering rather than generalized cytotoxicity. Together, these data indicate that **1a** selectively modulates a small subset of metastasis-associated transcripts without broad transcriptional reprogramming, supporting a mechanism primarily driven by post-translational modulation of *O-*glycosylation.

### 1a delays primary tumor growth and improves survival in a syngeneic mouse melanoma model

We investigated the therapeutic efficacy of GalNAc analogues *in vivo* using the syngeneic B16F10 melanoma model in C57BL/6J mice (31). We employed B16F10-Luc2 cells, which stably express firefly luciferase, enabling non-invasive, real-time monitoring of tumor growth via bioluminescence imaging (BLI), upon intraperitoneal administration of D-luciferin (32). We tested whether the intraperitoneally administered GalNAc analogues are systemically bioavailable and reach the subcutaneous tumor. To address this, we exploited metabolic glycan engineering (MGE) using Ac_4_GalNAz (**1e**), a peracetylated azide-bearing GalNAc analogue, followed by bio-orthogonal ligation with DBCO-Cy5 through strain-promoted azide-alkyne cycloaddition (SPAAC, also known as click chemistry) (33). After optimizing click reaction conditions *in vitro* (**Fig. 5 A**), C57BL/6J mice were implanted with B16F10-Luc2 cells subcutaneously (1.0 × 10^6^ cells per mouse, 50 µL) on day 1 and administered with either vehicle (DMSO, D) or **1e** (300 mg/kg, i.p.) on days 1, 3, 5, 7, and 9. On day 10, mice were euthanized, tumor masses were excised, and single-cell suspensions were prepared. Tumor cells were subjected to SPAAC with DBCO-Cy5, lysed, and resolved on an SDS-PAGE gel. Robust Cy5 fluorescence was detected exclusively in **1e**-treated samples, whereas vehicle-treated mice showed only a faint signal; silver-stained gels were used as loading controls (**Fig. 5 B**). These results showed that intraperitoneally administered GalNAc analogues are systemically bioavailable and efficiently incorporated into tumor cell glycoproteins *in vivo*, likely facilitated by the enhanced permeability and retention (EPR) effect characteristic of solid tumors.

**Figure 5.**
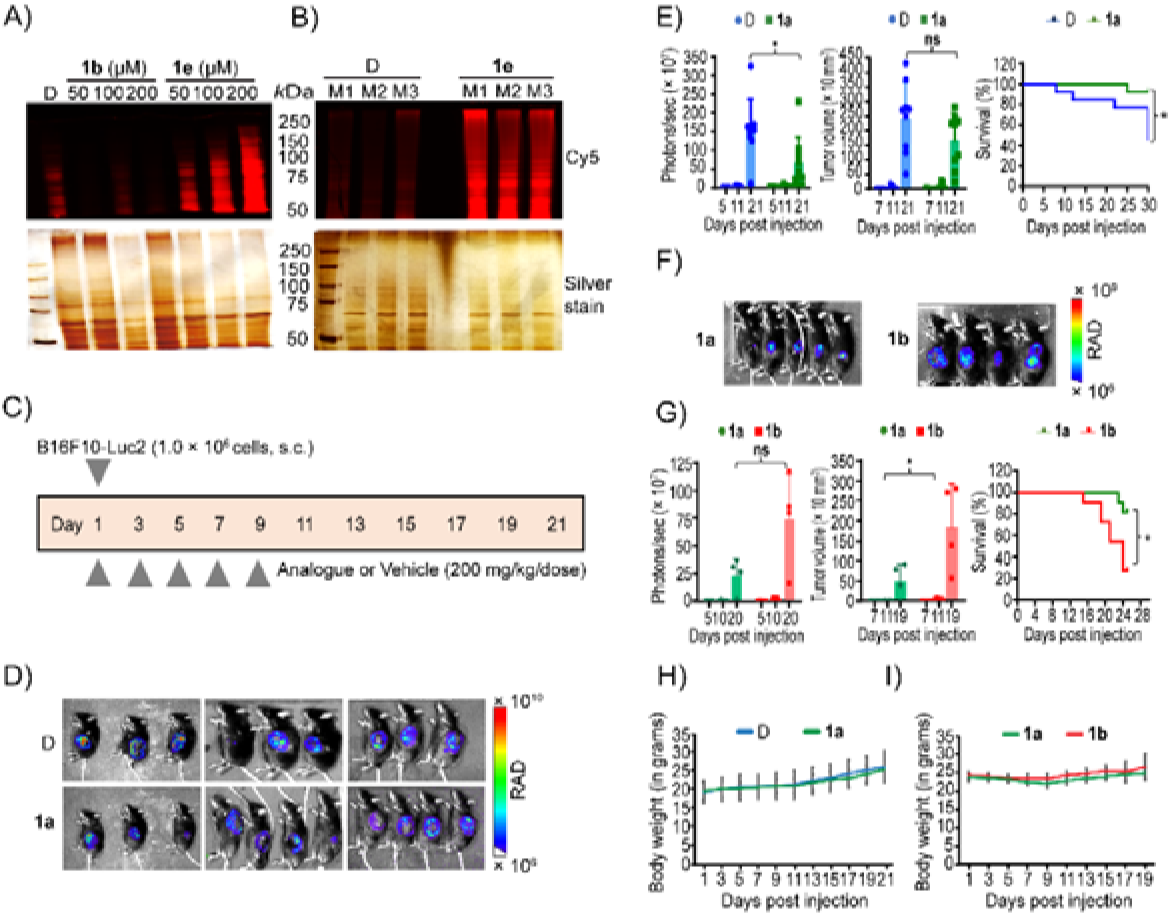
Tumor bioavailable-1a suppresses tumor growth and improves survival in a syngeneic melanoma mouse model. (A-B) Metabolic labeling confirms tumor bioavailability of intraperitoneally administered azido analogue (**1e**). (A) B16F10-Luc2 cells (1.0 × 10^6^) were treated with **1b** or **1e** at 50, 100, or 200 µM for 48 h, followed by reaction with DBCO-Cy5 (10 µM, 30 min) via copper-free SPAAC. Cells were lyzed using chloroform:methanol:water mixture, and proteins were precipitated with methanol. Protein samples (20 µL in SDS buffer) were resolved on 7.5% SDS-PAGE gels and imaged using a fluorescence scanner under the Cy5 (red) channel. Silver staining is shown as loading control. (B) *In vivo* bioavailability of **1e** at the tumor site. B16F10-Luc2 cells (1.0 × 10^6^ cells in 50 µL) were injected subcutaneously into the left flank of C57BL/6J male mice (6 weeks old) on day one, followed by i.p. (intraperitoneal) administration of **1e** (300 mg/kg) or D once on days one, three, five, seven, and nine. On day 10, tumors were excised, homogenized, and single cell suspensions were clicked with 10 µM DBCO-Cy5 for 30 min. Proteins were processed as described in (A) and imaged under the Cy5 channel (n = 3 mice per group). (C) Schematic representation of the experimental timeline and treatment regimen. (D) B16F10-Luc2 cells (1.0 × 10^6^ cells in 50 µL) were injected subcutaneously into the left flank of C57BL/6J male mice under anesthesia on day one, followed by i.p. administration of **1a** (200 mg/kg) or D once on days one, three, five, seven, and nine. Tumor progression was monitored by bioluminescence imaging (BLI) after i.p. injection of D-luciferin; a representative image from day 21 is shown. Two mice in the vehicle-treated group died on day 4 due to poor health unrelated to treatment. (E) Quantification of BLI signal (photons/sec) and tumor volume (mm^3^) measured using Aura software and vernier calipers, respectively, along with Kaplan-Meier survival analysis with an endpoint at day 30. Error bars represent the SD of n = 9 mice (three mice each in three independent experiments) for BLI/tumor volume and n = 13 for the survival curve. (F) B16F10-Luc2 cells (1.0 × 10^6^ cells in 50 µL) were injected subcutaneously on day one, followed by 200 mg/kg i.p. doses of **1a** or **1b** once, on days one, three, five, seven, and nine. One mouse in the **1b**-treated group died on day 18 due to tumor burden. Error bars represent the standard deviation of n = 5 for BLI/tumor volume and n = 11 for the survival curve. (G) Corresponding BLI (photons/sec), tumor volume quantification (mm^3^), and Kaplan-Meier survival curves with endpoints at day 25. Statistical significance was determined using multiple t-tests with the Holm-Sidak method (BLI and tumor volume) and the log-rank Mantel-Cox test (survival curve). P-value * ≤ 0.1 (BLI) and P-value * ≤ 0.05 (survival curve) in (E); P-value * ≤ 0.1 (tumor volume) and P-value * ≤ 0.01 (survival) in (G). (H-I) Body weight monitoring at every alternate day during treatment. (H) Comparison between vehicle (DMSO) and **1a**-treated groups; (I) Comparison between **1a**- and **1b**-treated groups. D, DMSO (vehicle); **1a**, Ac_5_GalNTGc; **1b**, Ac_4_GalNAc; and **1e,** Ac_4_GalNAz.

To evaluate therapeutic efficacy, B16F10-Luc2 cells (1.0 × 10^6^ cells) were implanted subcutaneously on day 1, followed by intraperitoneal administration of vehicle (D, 100 µL) or **1a** (200 mg/kg, 100 µL) on days 1, 3, 5, 7, and 9 (five injections in total) (**Fig. 5 C**). This study was performed across three independent cohorts with n = 3 mice per group per cohort (total n = 9 mice per group) to minimize inter-experimental variability in luciferase signal quantification arising from differences in luciferin administration and imaging conditions. Body weight was monitored on alternate days, tumor volume was measured on days 7, 11, and 21, and BLI was performed on days 5, 11, and 21. On day 21, representative BLI images revealed markedly decreased luminescence intensity in **1a**-treated mice compared with vehicle (D) control (**Fig. 5 D**). Quantitative analysis showed an approximately 60 % decrease in photon emission (**Fig. 5 E**, left panel) and a ∼40 % decrease in tumor volume compared to the vehicle control taken as 100 % (**Fig. 5 E**, middle panel), with no significant differences in body weight observed over the 21-day study period (**Fig. 5 H**). Kaplan-Meier survival analysis revealed 100 % survival (n = 13) in the **1a**-treated group through day 21, compared to 70 % survival in the vehicle group, attributable to the reduced tumor burden in the **1a**-treated mice (**Fig. 5 E**, right panel). The study was terminated on day 21 due to mortality observed in the vehicle-treated group. Remarkably, the tumor growth inhibition by **1a** persisted for 12 days after the final dose on day 9, demonstrating durable therapeutic efficacy without affecting body weight.

To validate the structure-activity relationships and confirm that the observed effects are specifically attributable to the thioglycolyl modification of **1a**, we compared **1a** with **1b** (wild-type control). Two groups (n = 5) of mice received B16F10-Luc2 cell implantation on day 1, followed by five intraperitoneal injections of either **1a** or **1b** on days 1, 3, 5, 7, and 9. On day 20, BLI images revealed a striking decrease in luminescence in the **1a**-treated cohort compared with **1b**-treated mice (**Fig. 5 F**). **1a**-treated mice exhibited a significant 60 % lower tumor burden, as measured by both photoluminescence (day 20; **Fig. 5 G**, left panel) and tumor volume (day 19; **Fig. 5 G**, middle panel), with no differences in body weight between groups (**Fig. 5 I**). Kaplan-Meier survival analysis again showed 100 % survival (n = 11) in the **1a**-treated group till day 21, compared to ∼70 % survival in the **1b**-treated cohort (**Fig. 5 G**, right panel).

Next, we benchmarked the efficacy of **1a** against doxorubicin (Dox), a standard anti-cancer agent (34). Dox was administered at a dose of 2.0 mg/kg (once weekly, for three weeks), a regimen reported to decrease tumor growth while minimizing systemic toxicity. Although Dox-treated mice showed greater tumor regression than the **1a** group, it was not statistically significant, while also exhibiting pronounced adverse effects, including decreased body weight, which were notably absent in **1a**-treated mice. Importantly, overall survival rates were comparable in both **1a** and Dox-treated groups, indicating that **1a** achieves therapeutic benefit similar to standard chemotherapy but with a superior safety profile (**Fig. 6, A-C)**. Finally, to confirm that the effects are strictly GalNAc-specific, we compared Ac_5_GalNAc (**1b**) with Ac_5_GlcNAc (**2b**), intraperitoneally. No significant differences in tumor volume were observed between mice treated with **1b** or **2b** (**Fig. 6, A, B,** and **D**). Together with the lack of efficacy of non-thiolated (**1b**) or non-GalNAc analogues (**2b**), these data confirm that the antitumor activity of **1a** is structure-dependent and requires both the GalNAc scaffold and the thioglycolyl modification.

**Figure 6.**
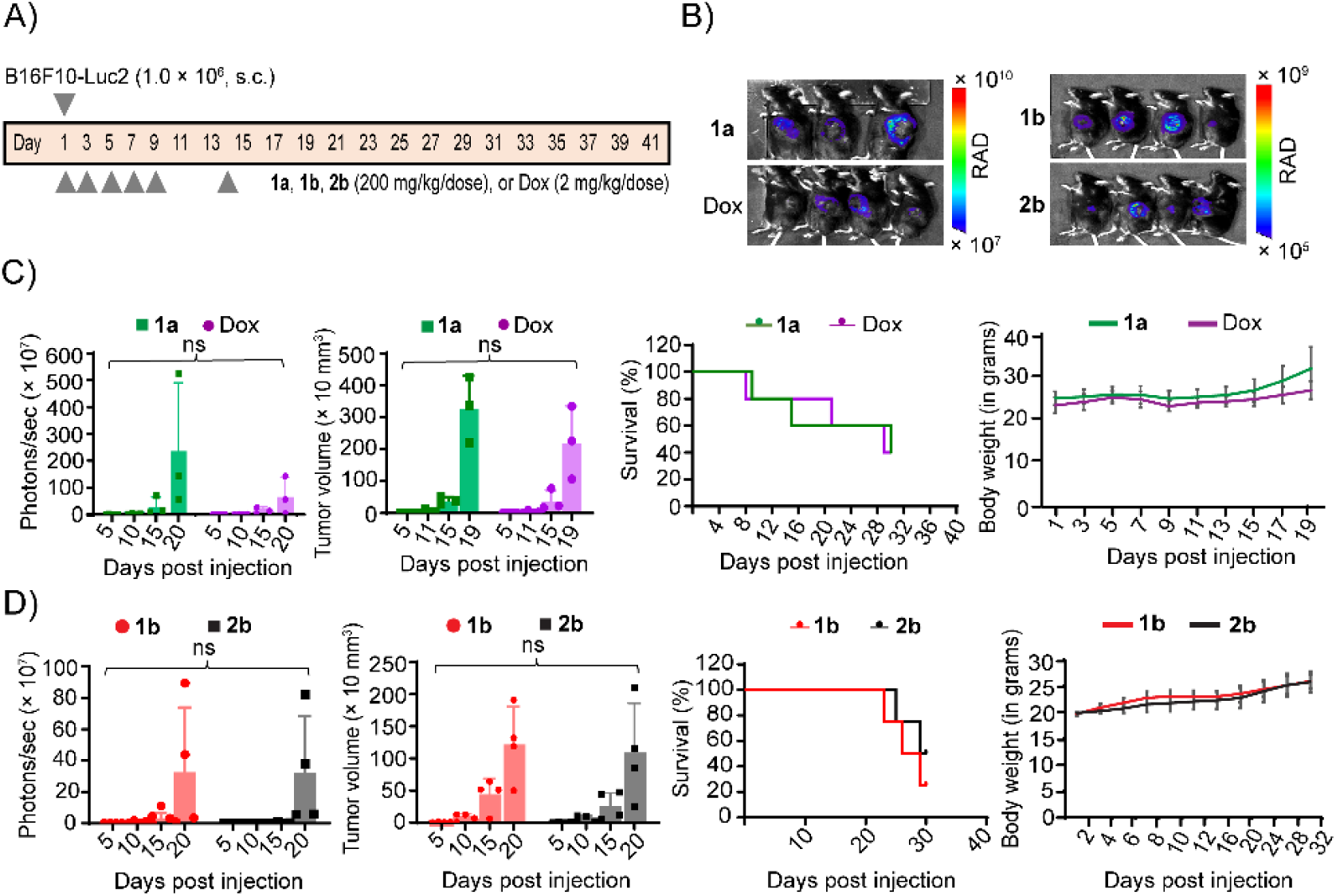
Benchmarking of 1a, and testing *in vivo* efficacy of 1b and its epimer 2b. (A) Schematic representation of the experimental timeline and treatment regimen. (B) Benchmarking of **1a** against Doxorubicin (Dox). B16F10-Luc2 cells (1.0 × 10^6^ cells in 50 µL) were injected subcutaneously into C57BL/6J male mice on day one. This was followed by i.p. administration of either **1a** (200 mg/kg) once on days one, three, five, seven, and nine, or Dox (2.0 mg/kg) once on days one, seven, and 14. Comparison between **1b** and its C-4 epimer **2b**. Mice bearing B16F10-Luc2 tumors were tested intraperitoneally with **1b** or **2b** (200 mg/kg) once on days one, three, five, seven, and nine. Representative images from day 20 are shown. (C-D) BLI was captured on days five, 10, 15, and 20 after D-luciferin administration. One mouse in the **1a-**treated group died on day 9 and day 15 each, and one mouse in the Dox-treated group died on day 9. Tumor burden (photons/sec), tumor volume (mm^3^), and body weight (grams) were quantified. Kaplan-Meier survival analysis was performed with an endpoint at day 30. Error bars represent the SD of n = 5 mice in all cases in (C) and n = 4 mice in (D). Statistical analysis was determined using multiple t-tests with the Holm-Sidak method (BLI and tumor volume) and the log-rank Mantel-Cox test (survival curve). All comparisons were non-significant. **1a**, Ac_5_GalNTGc; **1b**, Ac_4_GalNAc; and **2b**, Ac_4_GlcNAc.

Finally, safety profiling of **1a** in nontumor-bearing C57BL/6J mice, treated with various dosing regimens, was performed. It showed no changes in body weight following five daily doses of 200 mg/kg or 300 mg/kg, nor after ten daily doses of 200 mg/kg. Furthermore, there were no discernible alterations in blood counts, tissue pathology, or serum markers of hepatic or renal damage. Only mild kidney tubular dilatation was noted in the 200 mg/kg, 10-day prolonged dose group. Comparable findings were obtained in untreated, vehicle-treated, and **1b**- or **1c**-treated control groups (**Fig. S5 A**, **and Table 1**). Overall, **1a** administration is well-tolerated and does not result in significant systemic toxicity.

**Table 1.**
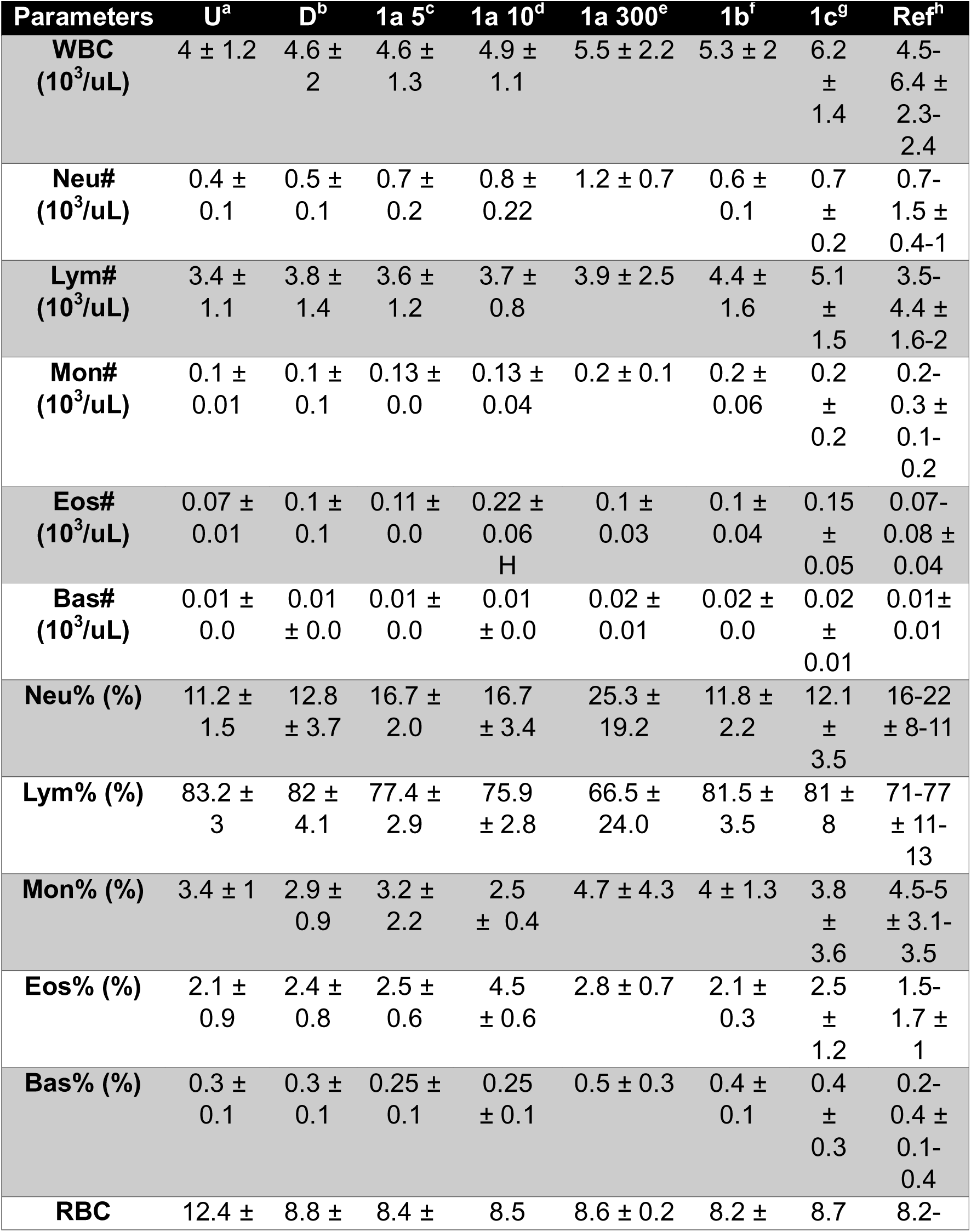
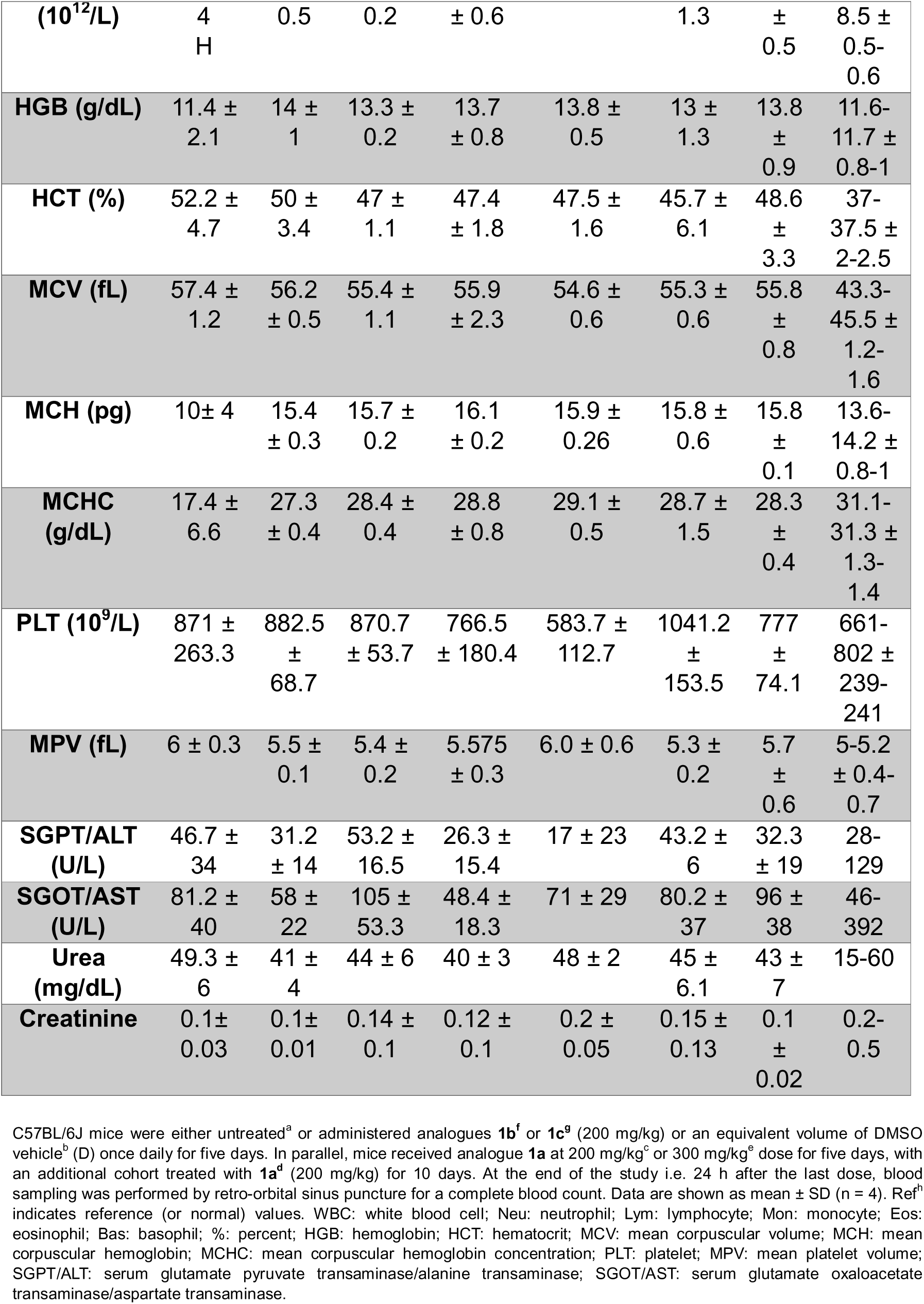
Hematological and serum biochemical assessment of HexNAc analogues in C57BL/6J mice.

### 1a pre-treated melanoma cells show severely impaired lung colonization in an experimental metastasis model

Having established that **1a** inhibits primary tumor growth and improves survival in orthotopic syngeneic melanoma models, we investigated whether MTOG inhibition affects the metastatic potential of melanoma cells *in vivo*. To directly assess the impact of **1a** on tumor cell extravasation and metastatic colonization, we employed an experimental lung metastases model (31). B16F10-Luc2 cells were pre-treated *in vitro* with vehicle (D, DMSO), wild-type control (**1b**), or **1a** (100 µM, 48 h), harvested, washed, and resuspended as single cell suspensions. Cells were then administered to C57BL/6J mice *via* retro-orbital intravenous injection (5.0 × 10^5^ cells in 50 µL per mouse) (**Fig. 7 A**). On day 19, mice were euthanized, and lungs were harvested.

**Figure 7.**
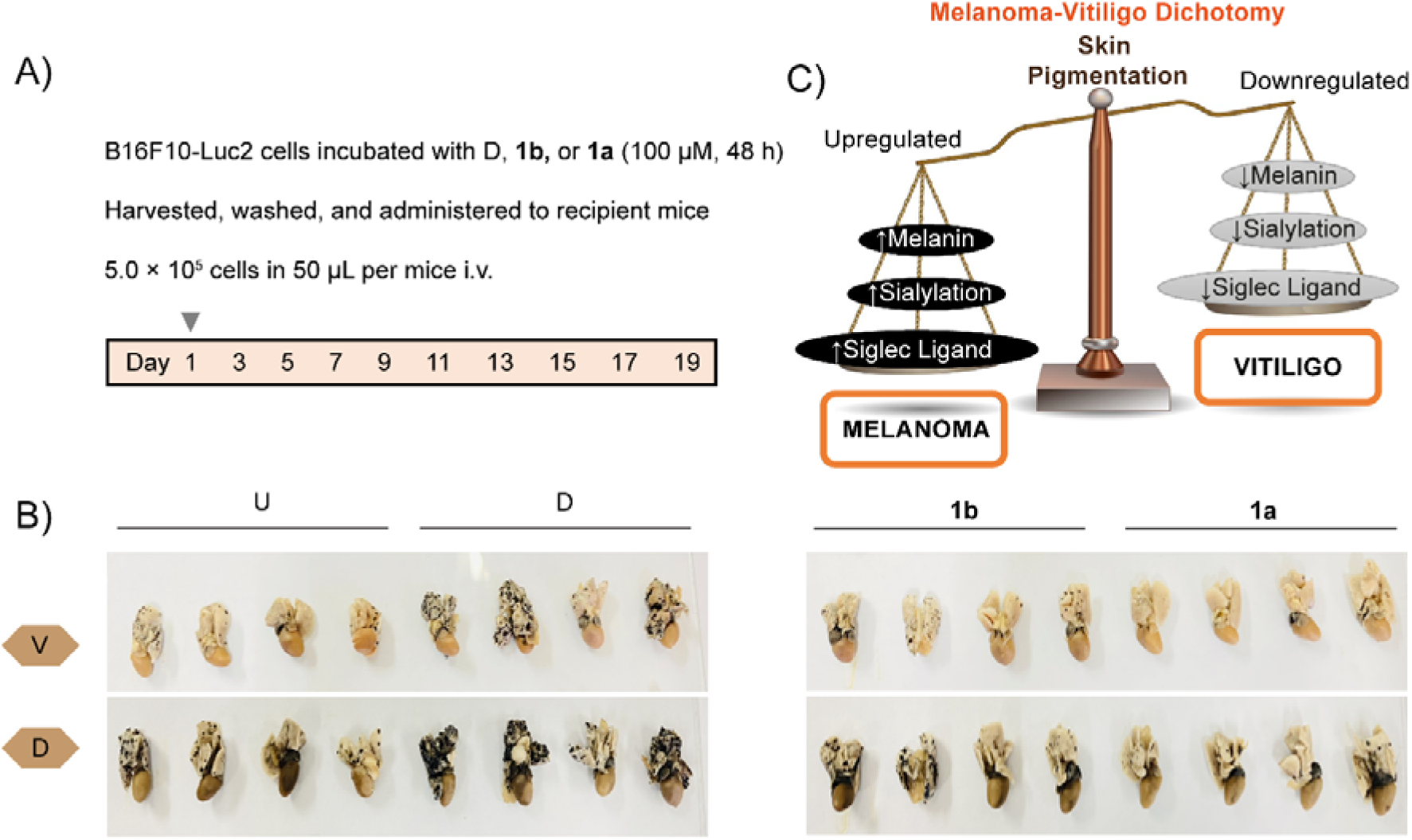
Intravenous injection of 1a-pretreated melanoma cells impairs metastatic lung colonization. (A) Schematic diagram showing the experimental timeline and treatment design. B16F10-Luc2 cells (5.0 × 10^5^) were either untreated (U) or treated with D, **1a**, or **1b** (100 µM each, 48 h). Cells were harvested by trypsinization, washed thoroughly, resuspended in PBS, and 5.0 × 10^5^ cells in 50 µL PBS were administered per mouse via the retro-orbital intravenous injection into C57BL/6J male mice (6 weeks old) under anesthesia. (B) Representative images of lungs (harvested together with the heart) collected on day 19 post-injection and imaged from both ventral (V) and dorsal (D) surfaces, showing metastatic melanoma burden. (C) Conceptual model illustrating the role of MTOG in regulating melanocyte and melanoma cell states. Balanced *O-*glycosylation supports normal pigmentation, whereas excessive melanin production and aberrant glycosylation contribute to melanoma progression and metastatic competence. Conversely, strong suppression of melanin synthesis is associated with hypopigmented states such as vitiligo and albinism. In the context of this study, **1a**-mediated inhibition of MTOG selectively decreases melanoma cells’ surface sialylation, associated Siglec-ligand presentation, and perturbs PMEL/gp100 glycosylation, thereby shifting melanoma cells away from a highly pigmented, immune-evasive, and metastatic phenotype without broadly altering gene expression programs. U, untreated; D, DMSO (vehicle); **1a**, Ac5GalNTGc; and **1b**, Ac4GalNAc.

Gross examination of harvested lungs revealed striking differences in metastatic burden among treatment groups. Control mice receiving either vehicle-treated or **1b**-treated B16F10-Luc2 cells developed numerous dark, pigmented melanoma nodules on both dorsal and ventral lung surfaces, consistent with robust pulmonary colonization by melanoma cells (**Fig. 7 B**). In sharp contrast, mice receiving **1a**-pre-treated melanoma cells showed a dramatic decrease in lung and cervical lymph node metastasis, with approximately 80 % fewer metastatic nodules than in **1b**-treated controls (**Fig. S5, B and C**). Importantly, this profound suppression of lung colonization occurred despite intravenous injection of identical numbers of tumor cells, indicating that **1a** treatment intrinsically compromises metastatic competence rather than cell viability. These findings establish that cell-surface sialylation and mucin-type *O-*glycan presentation are critical for melanoma cell survival in circulation, endothelial extravasation, and secondary site colonization (35).

Our working model suggests that **1a**-mediated inhibition of MTOG decreases melanoma surface sialylation and Siglec-ligand presentation, weakening glycan-dependent immune inhibitory interactions and compromising metastatic fitness (**Fig. 7 C and 1 A**). Concomitantly, decreased sialylation may enhance peptide-MHC (pMHC) surface presentation, density, or stability on tumor cells, potentially facilitating more efficient TCR engagement (36). Importantly, these effects arise without broad transcriptional reprogramming, highlighting the central role of post-translational glycosylation in melanoma progression.

## Discussion

This study identifies mucin-type *O-*glycosylation (MTOG) as a critical regulatory node in melanoma progression and a viable therapeutic target. By screening multiple GalNAc analogues, we identified Ac_5_GalNTGc (**1a**) as a lead compound that selectively inhibits *O-*glycan biosynthesis via metabolic incorporation. Unlike halogen-substituted analogues (like P-3Fax-Neu5Ac) that act through feedback inhibition, **1a** directly disrupts *O-*glycosylation, evidenced by a 75 % decrease in melanin and marked cell-surface hyposialylation (10). These results validate the thioglycolyl modification as a key determinant of efficacy.

Mechanistically, **1a** selectively disrupts PMEL17/gp100 glycosylation while sparing non-glycosylated proteins like MART-1, leaving the core transcriptional program intact. This confirms that **1a** acts post-translationally rather than through generalized cytotoxicity. The loss of HMB45 epitopes indicates that *O-*glycans play a structural role in melanosomal architecture, likely stabilizing amyloid fibrils by limiting aberrant protein-protein interactions and maintaining appropriate spacing between fibrils, thereby preserving matrix organization and structural integrity. In melanoma, where melanosome integrity is tightly coupled to pigmentation, immune suppression, and metastatic competence, **1a**-induced hypoglycosylation generates structurally defective melanosomes, blocking melanogenesis and attenuating malignant phenotypes.

While conventional depigmenting agents target tyrosinase activity, our findings reveal glycosylation as a more fundamental regulatory layer (37). This suggests that age-related or inflammatory glycan dysregulation may contribute to pigmentary disorders like vitiligo or lentigo. In vitiligo, glycan loss might destabilize melanosomes and expose cryptic antigens, precipitating immune-mediated melanocyte destruction.

The molecular mechanisms underlying melanoma progression are intricately linked to melanogenesis, melanin content, and pigmentation dynamics, encompassing both hyperpigmentation, such as lentigo, and depigmentation disorders, including vitiligo (representing two sides of the same coin). Conceptually, melanocyte biology can be viewed along a pigmentation continuum in which tightly regulated melanosome maturation and melanin production maintain normal skin pigmentation, whereas sustained, dysregulated melanogenesis supports melanoma progression, and impaired melanosome integrity or pigment synthesis leads to hypopigmentary disorders such as vitiligo (29) (**Fig. 7 C**). Importantly, this continuum reflects functional melanocyte output rather than changes in melanocytic gene expression per se. By selectively perturbing melanosomal *O-*glycosylation without altering transcriptional programs, **1a** shifts melanoma cells toward a hypopigmented, melanosome-defective state. This positions **1a**-treated cells as a valuable experimental model for interrogating glycosylation-dependent mechanisms governing melanosome stability, pigmentation, and immune recognition.

Our findings align with the growing evidence that melanogenesis and intact melanosome biology are closely linked to melanoma invasiveness and immune evasion (38). While melanin is photoprotective under physiological conditions, elevated melanin production in melanoma can promote oxidative DNA damage and support tumor evolution (3). In this context, selective attenuation of melanin synthesis through disruption of melanosomal *O-*glycosylation provides a mechanistically distinct strategy to impair both tumor-intrinsic and microenvironmental drivers of metastasis.

Treatment with **1a** markedly impaired melanoma cell migration and invasion *in vitro* and decreased Siglec-E binding by ∼35 %, thereby functionally disrupting the immunosuppressive Siglec-sialoglycan axis. Importantly, these molecular and cellular effects translated into robust therapeutic efficacy *in vivo.* Using a syngeneic mouse melanoma model, we observed that **1a** significantly delayed primary tumor growth upon repeated administration. This antitumor activity was likely facilitated by enhanced tumor accumulation of **1a**. Tumor bioavailability of intraperitoneally administered **1a** was experimentally validated using the azido-containing GalNAc analogue **1e** as a metabolic probe, confirming its incorporation within tumor tissues. Upon further evaluation, **1a** profoundly impaired lung colonization in an experimental metastasis setting and significantly improved overall survival. Notably, these antitumor effects were comparable to those achieved with doxorubicin, yet occurred without detectable systemic toxicity or weight loss, highlighting the favourable therapeutic index associated with of targeting MTOG.

In conclusion, this study demonstrates that pharmacological inhibition of MTOG by **1a**, exerts a dual anti-tumorigenic effect: (i) dismantling the melanosomal architecture essential for pigmentation, and (ii) disrupting Siglec-sialoglycan checkpoint-mediated immune evasion pathways. These findings highlight the largely untapped therapeutic potential of glycoengineering-based strategies in cancer treatment using small molecules with demonstrated antitumor activity in multiple tumor models. Although further medicinal chemistry optimization and pharmacokinetic refinement will be required, **1a** represents a first-in-class MTOG-targeting agent that could be strategically deployed as a lead scaffold in combination with kinase inhibitors or immune checkpoint blockade to overcome therapeutic resistance and improve clinical outcomes in advanced melanoma.

## Materials and Methods

### Chemical Synthesis and Reagents

Peracetylated non-natural GalNAc analogues **1a**, **1b, 1c, 1e, 2a,** and **2b** were synthesized according to previously described protocols (13,15,17–19).

### Cell Culture

B16F10 cells (a kind gift from Dr. Rajesh Gokhale, IML, NII) were cultured in sterile T-75 flasks or 100 mm tissue culture-treated dishes in DMEM supplemented with 10 % fetal bovine serum (FBS) and 1.0 % pen-strep (final concentration of 100 U/mL of penicillin and 0.1 mg/mL of streptomycin). Cultures were maintained in a humidified incubator at 37 °C with 5.0 % CO_2_. For B16F10-Luc2 cells (a kind gift from Dr. Santiswarup Singha, NBL, NII), which constitutively express luciferase, the selection antibiotic Blasticidin S hydrochloride was added to a final concentration of 10 µg/mL. Cell density was determined using a particle counter (Beckman-Coulter Z2) by suspending 100 µL of cell suspension in 10 mL PBS. Only cells in the logarithmic growth phase were used for experiments. Cells were detached using 0.25 % trypsin-EDTA, quenched with ∼10 mL of spent media, and pelleted by centrifugation (500 × g, 5.0 min). The supernatant was aspirated, and the pellet was resuspended in PBS and washed once by centrifugation (10 mL, 500 × g, 5.0 min). Hereafter, resuspending the cell pellet in PBS, ASB (avidin staining buffer), or LSB (lectin staining buffer), followed by centrifugation (500 × g, 5.0 min) to remove residual media, serum, antibodies, dye, or debris, is referred to as ‘one wash cycle’.

### Experiments *in vivo*

All animal protocols and experiments were approved by the Institutional Animal Ethics Committee (IAEC# 603/22) of the National Institute of Immunology (NII). Male C57BL/6J (WT) mice (4-6 weeks old) were procured from the Experimental Animal Facility (EAF), NII. Mice were provided food and water *ad libitum* and maintained under standard pathogen-free conditions. Parameters of weight, health, behaviour, and morbidity were monitored regularly. Mice showing adverse parameters were excluded from the study. Animals were anesthetized via intraperitoneal (i.p.) administration of 100 µL of ketamine (80 mg/kg) and xylazine (20 mg/kg) solution before cell implantation, imaging, or analogue injection. The anesthetic effect lasted for approximately 20-30 min, which was sufficient for all procedures. Animals were euthanized using CO_2_ asphyxiation whenever applicable. For i.p. injections, stock solutions of monosaccharide analogues and doxorubicin were prepared in DMSO and sterile filtered through 0.2 μm PTFE syringe filters. 100 µL of sterile-filtered DMSO served as the vehicle control.

### Estimation of Melanin Content

Melanin content was quantified according to an established protocol (39). B16F10 cells were seeded into 24-well plates (5.0 × 10^4^ cells/well in 500 µL complete medium) and treated with vehicle (D), **1a**, **1b**, **1c**, or **1d** (100 µM for 48 h). For dose-dependency studies, cells were treated with 25, 50, 100, or 200 µM of the respective analogues. After 48 h, the media containing floating cells was aspirated. Adherent cells were washed by gently swirling with 5.0 mL PBS, which was then aspirated. Cells were detached with 0.25 % trypsin (1.0 mL), neutralized with complete media (9.0 mL), and centrifuged at 500 × g for five min. The supernatant was discarded, and the cell pellet was dissolved in 100 µL of 1.0 N NaOH containing 1.0 % DMSO at 80 °C for 1 h. Subsequently, 100 µL of each lysate was transferred to a 96-well plate, and absorbance was measured at 405 nm using a microplate reader. Melanin concentration was determined using a standard curve generated from synthetic melanin (0 to 400 µg/mL) dissolved in 1.0 N NaOH. Phase-contrast images of adherent cells treated with D or **1a** were acquired at 10× magnification using an inverted Nikon microscope.

### Lectin staining for imaging by confocal microscopy

B16F10 cells were seeded at a cell density of 5.0 × 10^4^ cells/well in 500 µL complete medium in 24-well plates containing sterile 1.0 mm coverslips (12 mm diameter) and were either left untreated (U) or treated with vehicle (D), **1a**, **1b**, **1c**, or **1d** (100 μM, added to cell suspension before plating). After 48 h, the spent medium was aspirated, and the cells were gently washed with 0.5 mL PBS. Cells were then fixed with 0.5 mL of freshly prepared 4.0 % paraformaldehyde (PFA) in PBS for 10 min at room temperature (RT), followed by three washes with 0.5 mL of 0.1 % PBST (PBS containing 0.1% Tween 20). Cells were permeabilized with 0.5 mL of PBS containing 0.1 % Triton X-100 for 5 min at RT, then washed three times with 0.5 mL of 0.1 % PBST for 2.0 min each. Blocking was performed with 0.5 mL of 2.0 % gelatin in 0.1 % PBST for 30 min at RT, followed by six washes with 1.0 mL of 0.1 % PBST (2 min each). Cells were stained with fluorescein-conjugated wheat germ agglutinin (WGA; 1:2000 dilution from a 5.0 mg/mL stock) in 100 µL of 0.1 % PBST for 30 min at RT. Following lectin staining, nuclei were counterstained with DAPI (10 ng/mL in 100 μL PBS for 20 min at RT). Cells were then washed three times with 0.5 mL of 0.1 % PBST for 2.0 min each. One drop of ProLong Gold Anti-fade reagent was placed on a clean glass slide; after draining the residual PBST, coverslips were inverted and gently placed onto the antifade droplet to avoid air bubbles. Slides were cured overnight at 4 °C, and the edges were sealed with transparent nail polish prior to imaging. Samples were imaged on a Zeiss LSM 980 confocal microscope using a Plan-Apochromat 63× oil immersion objective. At least three coverslips (three biological replicates) were analyzed per condition, and three non-overlapping fields were acquired per coverslip. Fluorescence channels were assigned as follows: blue, DAPI; green, fluorescein-WGA.

### Lectin blotting for glycan profiling

B16F10 cells were treated with D, **1a**, **1b**, **1c**, or **1d** (100 μM, 48 h) while in suspension, alongside an untreated (U) control. After 48 h, the medium containing non-adherent cells was discarded. Adherent cells were detached with 0.25 % trypsin (1.0 mL), neutralized with complete medium (9.0 mL), and centrifuged at 500 × g for five min. The supernatant was aspirated, and the cell pellet was washed with PBS (3 × 5.0 mL at 3,000 × g for 2.0 min each). Cell lysates were prepared in 100 µL RIPA buffer supplemented with PIC (1:100). Protein concentration was determined using the Bradford assay. Equal amounts of protein were resolved on 7.5 % SDS-PAGE and transferred to nitrocellulose membranes at a constant current of 250 mA for three h at 4 °C. Membranes were blocked for 1 h at RT with 2.0 % gelatin in 0.1 % PBST (10 mL per membrane). Blocked membranes were incubated for 1 h with biotinylated *Maackia amurensis* lectin-II (MAL-II; 1:2000 from a 1.0 mg/mL stock) or fluorescein-conjugated *Vicia villosa* agglutinin (VVA; 1:2000 from a 2.0 mg/mL stock) diluted in 5.0 mL of 0.1 % PBST. Membranes were then washed six times with 10 mL of 0.1 % PBST for five min each. VVA blots were visualized on an Amersham Typhoon imager using the Cy2 channel. For MAL-II detection, membranes were incubated for 30 min with FITC-avidin (1:50,000 from a 2.0 mg/mL stock) in 5.0 mL of 0.1 % PBST, followed by six additional washes with 10 mL of 0.1 % PBST (five min each). MAL-II blots were then imaged on the Amersham Typhoon using the Cy2 filter. Three biological replicates were performed for each lectin.

### Lectin flow cytometry

Lectin binding was assessed using a reported protocol (13). B16F10-Luc2 cells were harvested by standard trypsinization as described above. For treatment, 1.5 mL of complete medium was added to each well of a six-well plate, and wells were left untreated (U) or treated with vehicle (DMSO), **1a**, **1b**, **1c,** or **1d** (100 µM final concentration). Cells (3.0 × 10^5^ cells in 1.5 mL complete medium; final density 1.0 × 10^5^ cells/mL) were added to each well and incubated for 48 h at 37 °C. After treatment, cells were detached with trypsin and collected by centrifugation (500 × g, 5.0 min). Supernatants were discarded, and pellets were resuspended in 5.0 mL complete medium and incubated at 37 °C for 30 min to allow replenishment of cell surface glycans. Cells were then pelleted again (500 × g, 5.0 min) and resuspended in 1.0 mL PBS to obtain a density of 2.0 × 10^5^ cells/mL. Suspensions were transferred to pre-labelled microcentrifuge tubes (MCTs) and washed twice with 1.0 mL PBS. For neuraminidase controls, 2.0 × 10^5^ cells were resuspended in 100 μL PBS and incubated with Neuraminidase from *Vibrio cholerae* (final concentration of 20 milliunits/mL) at 37 °C for 30 min, followed by two PBS washes (1.0 mL each). For lectin staining, cells were resuspended in either 500 µL lectin staining buffer (LSB) containing SNA (0.5 µg/mL) or 200 µL LSB containing VVA (10 µg/mL) and incubated for 15 min at 4.0 °C. Cells were washed once with 500 µL LSB, then resuspended in 200 µL LSB, and 0.5 µL propidium iodide (2.5 µg/mL) was added to each sample immediately before acquisition. Flow cytometry data were acquired by sequential gating: FSC-A/SSC-A to exclude debris, FSC-H/FSC-A to exclude doublets, followed by gating on PI-A-negative live cells; 10,000 live events were collected per sample. Two biological replicates per condition, each with two technical replicates, were analyzed using FlowJo v10, and geometric mean fluorescence intensity (GMFI) of FITC-positive populations was plotted.

### Immunoprecipitation and Lectin Blotting of PMEL17/gp100

Untreated and **1a**-treated (100 µM, 48 h) B16F10 cells were lysed in RIPA buffer as described previously, and total protein concentration was determined using the Bradford assay. For each immunoprecipitation reaction, 1 mg of total cell lysate was incubated with 1 µg of anti-gp00 (*N-*term) antibody overnight at 4 °C with end-to-end rotation to facilitate antigen-antibody binding. Subsequently, 50 µL of pre-washed Protein G agarose beads (equivalent to 20 µL of bead slurry, Merck; 16-266) were added to the immune complexes. The mixture was incubated for 2 h at 4 °C, followed by an additional 1 h at RT on an end-over-end rotor. Bead-antibody-antigen complexes were pelleted by centrifugation at 1400 rpm for 5 min, and the supernatant was collected as the ‘depleted’ fraction. The beads were washed twice with 250 µL of 0.1 % PBST using gentle wave motion mixing. Immunoprecipitated proteins were eluted twice with 30 µL of 0.2 M Glycine (pH 2.6). Aliquots from all fractions, input, depleted, wash, elution, and boiled beads (BB), were boiled at 95 °C for 10 min. Samples were resolved by SDS-PAGE on a 10 % polyacrylamide gel at 120 V and transferred to a nitrocellulose membrane as described above.

For PMEL17/gp100 detection, the membrane was blocked for one h with 5.0 % (w/v) non-fat milk (NFM) (5.0 mL) in PBS. The blocked membrane was incubated overnight at 4 °C with anti-gp100 (*N*-term) primary antibody (1:10,000 dilution, stock conc. 0.357 mg/mL) in 5.0 mL of 5.0 % (w/v) NFM in PBS. Membrane was washed thrice with 0.1 % PBST (10 mL/wash, 5 min each) and incubated for 1 h at RT with AF488-conjugated anti-rabbit IgG secondary antibody (1:10,000 dilution, stock conc. 2.0 mg/mL) in 5.0 mL 0.1 % PBST. Following three additional washes with 0.1 % PBST (10 mL/wash, 5 min each), signals were visualized using an Amersham Typhoon fluorescent scanner. Glycan analysis was performed using lectin blotting with MAL-II and VVA as described previously. Two biological replicates, each with two technical replicates, were performed for each antibody or lectin probe.

### Western Blotting for Melanosomal Proteins

Cell lysates from analogue-treated samples were prepared as described above. Proteins were resolved by 7.5 % SDS-PAGE and transferred to a nitrocellulose membrane using a wet transfer system at a constant current of 250 mA for three h at 4 °C. Membranes were blocked for one h at RT with 5.0 % (w/v) NFM (5.0 mL) in PBS. Following blocking, membranes were incubated overnight at 4 °C with the following primary antibodies diluted in 5.0 mL of 5 % (w/v) NFM in PBS: anti-gp100 (*N*-term; 1:10,000, stock conc. 0.357 mg/mL); anti-gp100 (HMB45; 1:1000, stock conc. 0.2 mg/mL); anti-gp100 (C-term; 1:1000, stock conc. 0.5 mg/mL); anti-tyrosinase (1:1000, stock conc. 0.2 mg/mL); anti-MART-1 (1:5000, stock conc 1.08 mg/mL); and anti-β-actin (1:20,000, stock conc. 2.0-2.5 mg/mL). Membranes were washed thrice with 0.1 % PBST (10 mL) for five min each. Subsequently, membranes were incubated for 1 h at RT with the corresponding 2° antibodies (diluted 1:10,000 in 5.0 mL of 0.1 % PBST). AF488-conjugated anti-rabbit IgG (stock conc. 2.0 mg/mL) was used for gp100 (*N*-term), tyrosinase, and MART-1 detection. Cy5-conjugated donkey anti-mouse IgG (stock conc. 1.4 mg/mL) was used for gp100 (HMB45) and β-actin, while AF488-conjugated anti-mouse IgG (stock conc. 2.0 mg/mL) was used for gp100 (*C*-term). After three final washes with 0.1 % PBST (10 mL, five min each), fluorescent signals were visualized using an Amersham Typhoon scanner. Experiments were conducted with two biological replicates and two technical replicates per antibody probe.

For the dose dependency study of gp100 (*N-*term) and gp100 (HMB45), cells were either left untreated (U) or treated with vehicle (D), **1a**, **1b**, **1c**, or **1d** at concentrations of 25, 50, 100, and 200 µM. For the time-course study, B16F10 cells were either left untreated (U) or treated with vehicle (D), **1a**, **1b**, **1c**, or **1d** (100 μM) for various time points (24, 48, 72, and 96 h). The remainder of the protocol was performed as described above. Two biological replicates, each with two technical replicates, were performed for this experiment.

### Staining of PMEL17/gp100 for Confocal Microscopy

B16F10 cells were analogue-treated, fixed, and permeabilized as described previously. The cells were washed three times with 0.5 mL of 0.1 % PBST for 2.0 min each. For F-actin staining, cells were incubated with 66 nM rhodamine phalloidin (0.2 µL; stock: 66 µM in DMSO) in 200 µL of 3.0 % BSA in PBS for 20 min at RT. Following staining, cells were washed thrice with 0.5 mL of 0.1 % PBST for two min each. Cells were incubated with primary antibodies in 0.1 mL of PBS overnight at 4 °C. The following primary antibodies were used: unconjugated anti-gp100 (*N*-term) (dilution 1:100, stock conc. 0.357 mg/mL) or anti-gp100 (HMB45) (dilution 1:100, stock conc. 0.2 mg/mL). The cells were washed thrice with 0.5 mL of 0.1 % PBST for five min each. Subsequently, cells were incubated with the corresponding secondary antibodies in 0.1 mL of PBS for 2 h at RT in the dark: Alexa Fluor 488 anti-rabbit IgG (dilution 1:1,000, stock conc. 2.0 mg/mL) for gp100 (*N*-term) or Cy5-conjugated donkey anti-mouse IgG (dilution 1:1,000, stock conc. 1.4 mg/mL) for gp100 (HMB45). After staining, cells were washed three times with 0.5 mL of 0.1 % PBST for 2.0 min each. Following antibody staining, cells were incubated with DAPI (10 ng/mL) in 100 µL of PBS for 20 min at RT. Cells were washed three times with 0.5 mL of 0.1 % PBST for 2.0 min each. Slides were mounted and visualized as described previously. At least three coverslips, representing three biological replicates, were scanned. For each biological replicate, three fields were imaged per coverslip. Fluorescence channels were assigned as follows: DAPI (Blue), Rhodamine Phalloidin (Red), Alexa Fluor 488 for PMEL17/gp100 (*N*-term) (Green), and Cy5 for PMEL/gp100 (HMB45) (Red).

### Colocalization of Melanosomal Proteins by Confocal Microscopy

B16F10 cells were **1a**-treated, fixed, and permeabilized as described previously. Cells were incubated with unconjugated anti-MART-1 (clone 48-118) antibody (dilution 1:100, stock conc. 1.04 mg/mL) in 0.1 mL of PBS overnight at 4 °C. The cells were washed thrice with 0.5 mL of 0.1 % PBST for five min each. Subsequently, cells were stained with Alexa Fluor 488 anti-rabbit IgG (dilution 1:1,000, stock conc. 2.0 mg/mL) in 0.1 mL of PBS for 2 h at RT in the dark. The cells were then washed thrice with 0.5 mL of 0.1 % PBST for two min each. Following MART-1 staining, cells were incubated with conjugated Alexa Fluor 647 anti-gp100 (*N-*term) antibody (dilution 1:100, stock 0.5 mg/mL) in 0.1 mL of PBS overnight at 4 °C. The cells were washed thrice with 0.5 mL of 0.1 % PBST for five min each. For nuclear counterstaining, DAPI (10 ng/mL) was added in 100 µL of PBS and incubated for 20 min at RT. Cells were washed three times with 0.5 mL of 0.1 % PBST for two min each. Slides were mounted and visualized as described previously. At least three coverslips, representing three biological replicates, were scanned. For each biological replicate, three fields were imaged per coverslip. DAPI (Blue), Alexa Fluor 488 for MART-1 (Green), and Alexa Fluor 647 for PMEL17/gp100 (*N-*term) (Red). Yellow signals indicate the colocalization of MART-1 with PMEL17/gp100.

### Quantification of Siglec-E-Fc Binding by Flow Cytometry

B16F10-Luc2 cells were harvested using standard trypsinization protocols as previously described. Cells (1.0 × 10^6^) were seeded in 5.0 mL of complete media (final cell density of 1.0 × 10^5^ per mL) into 10-cm tissue-culture treated plates. Plates were either left untreated or treated with vehicle DMSO (D), **1a** (100 µM), **1b** (100 µM), **2a** (100 µM), or Dox (100nM) and incubated for 48 h. Following incubation, cells were trypsinized, harvested, and washed twice with PBS (10 mL). Aliquots of 1.0 × 10^6^ cells were prepared for analysis. For the negative control, 1.0 × 10^6^ cells were resuspended in 100 μL of PBS in an MCT and incubated with Neuraminidase from *Vibrio cholerae* (final concentration of 20 milliunits/mL) at 37 °C for 1 h. Cells were subsequently washed twice with PBS (1.0 mL). A 50 µL antibody pre-complex was prepared by mixing recombinant Siglec-E-Fc chimera (final concentration of 75 nM) and Alexa Fluor 488-conjugated goat anti-mouse IgG secondary antibody (final concentration of 50 nM) in PBS in an MCT. The pre-complex was incubated at 4 °C for 30 min. This pre-complex was then added to the cell aliquots and incubated for 30 min at 4 °C. A secondary antibody-only control was included to account for non-specific binding. Cells were washed twice with FACS buffer (1.0 mL), resuspended in 300 μL of FACS buffer, and analyzed. Cells were gated sequentially on FSC-A/SSC-A (to exclude debris), and FSC-H/FSC-A (for doublet exclusion). For each sample, 10,000 events were acquired. Two biological replicates (BR) were prepared, with two technical replicates (two counts per sample) for each BR. Siglec-E-Fc binding was analyzed using FlowJo software, and the MFI was plotted for the AF488-positive population.

### Migration and Invasion Assays

Migration assays were performed following a reported protocol (40). B16F10-Luc2 cells were harvested using standard trypsinization protocols as previously described. The resulting cell pellet was washed twice with PBS (10 mL). 1.0 × 10^5^ cells were resuspended in complete media (0.3 mL) and seeded into transwell inserts equipped with an 8.0 µm pore size PET membrane for migration. For invasion, the transwell inserts were pre-coated with Matrigel, mimicking the extracellular matrix. The inserts were carefully placed into a 24-well plate using forceps, establishing a two-chamber system: the upper chamber (insert) containing the cells (0.3 mL), and the bottom chamber (well) filled with complete media (0.7 mL) as a chemoattractant. Cells in the upper chamber were either left untreated (U) or treated with vehicle (D), **1a** (100 µM), or **1b** (100 µM) in a total volume of 1.0 mL. After cells adhered to the insert membrane (∼ 12 h), the complete media in the upper chamber was replaced with incomplete media (without FBS). Simultaneously, the media in the bottom chamber was replaced with fresh complete media to establish a chemotactic gradient. Cells were then either left untreated (U) or replenished with vehicle (D), **1a** (100 µM), or **1b** (100 µM) in a total volume of 1.0 mL and incubated for 48 h. Media was aspirated from the upper and bottom chambers, and the inserts were washed once by dipping them in a 100 mL beaker filled with PBS. The inserts were then placed into fresh wells, and 0.5 mL of 0.25 % aq. crystal violet stain was added to both the upper and bottom chambers. After a 20 min incubation, the inserts were washed twice by dipping them into a PBS-filled 100 mL beaker. Non-migrated cells on the interior of the upper chamber were removed using a cotton-tipped pipette without damaging the membrane. The upper chambers were placed directly on an inverted bright-field microscope, and images were acquired using a 10× objective (Nikon Digital Sight 10). Subsequently, the membranes were excised from the inserts using a surgical blade and placed into a 96-well plate. The membranes were soaked in 100 µL of DMSO to solubilize the stained cells, and absorbance was measured at 570 nm. Three biological replicates were prepared, with two technical replicates for each BR.

For the dose dependency study, inserts were either left untreated (U) or treated with vehicle (D), **1a**, or **1b** at 50, 100, and 200 µM concentrations. The remainder of the protocol was performed as described above. Two biological replicates, each with two technical replicates, were prepared for this experiment.

### Metabolic glycan engineering (MGE) to Test Tumor Accessibility to GalNAc Analogues

Optimization of MGE conditions in melanoma cells was performed following a reported protocol prior to *in vivo* testing (18). B16F10-Luc2 cells were analogue-treated for 48 h and harvested using standard trypsinization as previously described. At the indicated time point, cells were counted, and aliquots of 1.0 × 10^6^ cells were pelleted by centrifugation (500 × g, 5.0 min). The resulting pellets were washed with PBS (10 mL). Cell pellets were resuspended in 200 µL of PBS and subjected to a SPAAC reaction with 10 μM DBCO-Cy5 for 30 min. Two technical replicates were performed for each condition. Following the reaction, cells were immediately lysed by adding 500 µL of a chloroform:methanol:water (1:4:3) mixture. Proteins remaining in the solution were precipitated using methanol to yield a protein pellet. This pellet was resuspended in methanol and washed three times (centrifugation at 16000 × g, 5 min). Finally, the protein pellets were resuspended in 100 µL of 1.0 % SDS buffer. Protein concentration was determined using the Bradford assay. Approximately 20 µg of protein was loaded onto 7.5 % SDS-PAGE gels and resolved until the tracking dye reached the bottom of the gel. The gels were washed once with PBS (10 mL) in a glass container. Subsequently, the gels were placed in an Amersham Typhoon fluorescence imager and directly visualized using the Cy5 (red) channel. Silver-stained gels, run in parallel, served as loading controls. Two biological replicates, each with two technical replicates, were performed for each condition.

*In vivo* metabolic engineering using **1e** was assessed following a previously described protocol (33). Mice received intraperitoneal injections of either 300 mg/kg of **1e** or vehicle (D) once on days one, three, five, seven, and nine. On day 10, mice were euthanized, and tumors were excised and homogenized. Cell suspensions were prepared and processed for protein extraction and SPAAC labelling as described above. Three biological replicates, derived from three distinct mouse samples (numbered 1, 2, and 3), were analyzed with two technical replicates for each.

### Implantation of Melanoma Cells Followed by Intraperitoneal HexNAc Analogue Administration

B16F10-Luc2 cells were harvested using standard trypsinization protocols as previously described. Cells were counted, and aliquots of 1.0 × 10^6^ cells were pelleted by centrifugation (500 × g, 5 min). Pellets were washed twice with PBS (10 mL). For implantation, cells (1.0 × 10^6^) were resuspended in 50 µL of phosphate-buffered saline (PBS) and injected subcutaneously using a 27 ½-inch gauge syringe needle.

#### DMSO vs 1a

Following subcutaneous tumor cell implantation, mice (n = 9; three per group in three independent experiments) received intraperitoneal injections of either 100 µL of vehicle DMSO (D) or **1a** (200 mg/kg). Injections were administered once on days one, three, five, seven, and nine post-implantation. Analogues were administered under anesthesia whenever the dosing day coincided with an imaging session.

#### 1a vs 1b

Following subcutaneous tumor cell implantation, mice (five per group) received intraperitoneal injections of either 100 µL of **1a** (200 mg/kg) or **1b** (200 mg/kg). Injections were administered once on days one, three, five, seven, and nine post-implantation. Analogues were administered under anesthesia whenever the dosing day coincided with an imaging session.

#### 1a vs Dox

Following subcutaneous tumor cell implantation, mice (five per group) received intraperitoneal injections of either 100 µL of **1a** (200 mg/kg) or Dox (2.0 mg/kg). **1a** was administered once, on days one, three, five, seven, and nine under anesthesia. **Dox** was administered once on days one, seven, and 14. Analogues were administered under anesthesia whenever the dosing day coincided with an imaging session.

#### 1b vs 2b

Following subcutaneous tumor cell implantation, mice (four per group) received intraperitoneal injections of either 100 µL of **1b** (200 mg/kg) or **2b** (200 mg/kg). Injections were administered once on days one, three, five, seven, and nine post-implantation. Analogues were administered under anesthesia whenever the dosing day coincided with an imaging session.

### Tumor Size and Volume Measurement

Tumor growth was monitored in real-time using BLI following treatment with the substrate D-luciferin (stock concentration of 15 mg/mL in PBS). Mice were anesthetized and injected i.p. with 150 µL of D-luciferin (1.5 mg per mouse). Imaging was performed using an *in vivo* imaging system (Lago X). To prevent signal saturation, the exposure time was fixed at 60 s, and binning was adjusted from low to medium. Bioluminescence intensity was quantified in photons/sec using Aura software (version 2.03).

Tumor dimensions were measured at regular intervals using digital vernier calipers. Tumor volume was calculated using the formula [tumor volume in mm^3^ = (length × width^2^)/2]. Body weight was measured and recorded every alternate day, beginning on day one and continuing until the end of the study.

### Intravenous Administration of B16F10-Luc2 Cells

B16F10-Luc2 cells were harvested using standard trypsinization protocols as previously described. Cells (1.0 × 10^6^) were seeded in 5.0 mL of complete media (final cell density of 1.0 × 10^5^ per mL) into 10-cm tissue-culture treated plates containing 5.0 mL of complete media. Plates were either left untreated (U) or treated with vehicle (D), **1a** (100 µM), or **1b** (100 µM) and incubated for 48 h. Following incubation, cells were detached by trypsinization and washed twice with PBS (10 mL). Cells were counted, and aliquots of 5.0 × 10^5^ cells were pelleted by centrifugation (500 × g, 5.0 min). The final cell pellet was resuspended in 50 µL of PBS. The cell suspension was administered via the retro-orbital plexus using a 27½ inch-gauge syringe needle. Injections were performed at an acute angle with the bevel facing up at a slow rate to minimize injury. Four mice were included per group for this study.

### Harvesting, Bioluminescence Imaging, and Fixation of Lungs

Mice were euthanized via CO_2_ asphyxiation on day 19. The lungs and heart were dissected *en bloc* and immersed in sterile PBS (10 mL) to remove residual blood. For imaging, the lungs were placed in a 3.5-cm diameter dish and incubated with 100 µL of D-luciferin (15 mg/mL). BLI was acquired with an exposure time of 20 s and a medium binning setting. Following imaging, the lungs were rinsed in sterile PBS to remove residual D-luciferin. The tissues were then stored in glass vials containing 10 mL of Fekete’s solution (prepared using 100 mL of 70 % alcohol, 10 mL of 10 % formalin, and 5.0 mL of glacial acetic acid) (41). Under these conditions, melanoma nodules appeared black against the pale white background of the lung tissue. Nodules were also observed in the cervical lymph nodes. High-definition images were captured using a 64-megapixel smartphone camera.

### MTT assay for cell viability

This assay measures cellular metabolic activity as an indicator of cytotoxicity (42). It is a colorimetric assay based on the reduction of the yellow tetrazolium salt (3-(4,5-dimethylthiazol-2-yl)-2,5-diphenyltetrazolium bromide (MTT), to purple formazan crystals by metabolically active cells. Viable cells contain NAD(P)H-dependent oxidoreductase enzymes that catalyze this reduction. The insoluble formazan crystals are dissolved, and the resulting-colored solution was quantified by measuring absorbance. B16F10 cells were seeded in 96-well flat bottom plates (5.0 × 10^4^ cells per mL) in 200 μL of complete medium with phenol red (10,000 cells/well). Cells were either left untreated (U), or treated with vehicle (D), **1a, 1b, 1c,** or **1d** while in suspension at concentrations of 25, 50, 100, and 200 μM for 48 h at 37 °C. After 48 h, 10 μL/well of MTT labelling reagent (stock solution: 5.0 mg/mL in water; final concentration 0.5 mg/mL) was added to each well. The cells were incubated with MTT for 4 h at 37 °C, after which purple formazan crystals were visually confirmed under the microscope. The media was aspirated from all wells, and 100 μL/well of DMSO was added to dissolve the formazan crystals. The crystals were resuspended thoroughly using a multi-channel pipette, and absorbance was measured at 570 nm using a microplate reader.

### Trypan Blue Staining Assay

Trypan blue staining is a widely used method for measuring cell viability (43). Trypan blue is an azo dye impermeable to intact cell membranes; therefore, it enters only cells with compromised membranes (dead cells). Upon entry, the dye binds to intracellular proteins, staining the cells a distinctive blue colour. This allows for the differentiation of live (unstained) and dead (blue) cells within a population. B16F10 cells were seeded in a 24-well plate (5.0 × 10^4^ cells per mL/well) in 500 μL of complete medium (25k cells/well). Cells were either left untreated (U) or treated with vehicle (D), **1a, 1b, 1c,** or **1d** (100 μM, 48 h) while in suspension. After 48 h, the media containing floating cells was collected into fresh MCT. Adherent cells were detached by incubation with 0.25 % trypsin (500 μL), followed by dilution with complete media (2.0 mL). The floating cell fraction and the trypsinized cell suspension were combined and centrifuged at 500 × g for five min. The resulting cell pellet was resuspended in 1.0 mL of PBS. For counting, 5.0 μL of 0.4 % (w/v) trypan blue (40 mg trypan blue powder in 10 mL PBS) was mixed with 5.0 μL of the cell suspension and incubated for three min at RT. A 10 μL aliquot of the mixture was loaded onto a hemocytometer. Cells were counted in the four large corner squares and the central square under a microscope. Viable (unstained) and non-viable (stained) cells were counted separately. The total cell count across five squares was approximately 120 ± 15 cells. The total number of viable cells per mL was calculated by multiplying the total number of viable cells counted by the dilution factor. The total cell count per mL was determined by summing the viable and non-viable cells from the five squares. Cell viability was calculated using the following formula: Viable cells (%) = (total number of viable cells/total number of cells) × 100.

### Assay for cell proliferation

The stable incorporation of the intracellular fluorescent dye 5-(and -6)-carboxyfluorescein diacetate succinimidyl ester (CFSE) monitors cell division, as fluorescent labelling decreases sequentially in daughter cells (44). B16F10 cells at 0 h were resuspended in PBS containing 5.0 % FBS at a concentration of 5.0 × 10^4^ cells/mL. To label the cells, 1.0 μL of CFSE solution (stock 5.0 mM; 2.8 mg CFSE in 1.0 mL DMSO) was added to 1.0 mL of the cell suspension to achieve a final concentration of 5.0 μM, followed by rapid mixing. After a five min incubation at RT, 10 mL of PBS containing 5.0% FBS was added to quench the reaction. Cells were centrifuged for five min at 300 × g at 20 °C, and the supernatant was aspirated. The cells were washed thrice by resuspending in 1.0 mL of PBS containing 5.0 % FBS. Following staining, B16F10 cells were seeded in 100 mm diameter tissue-culture treated dishes (5.0 × 10^4^ cells/mL) in 10 mL of complete medium. Cells were either left untreated (U) or treated with vehicle (D), **1a, 1b, 1c,** or **1d** (100 μM, 48 h) while in suspension. After 48 h, media containing any floating cells was aspirated. Adherent cells were washed by adding PBS (5.0 mL), swirling gently, and aspirating. Cells were detached by incubating with 0.25 % trypsin (1.0 mL), followed by dilution with complete media (9.0 mL). The cell suspension was centrifuged at 500 × g for five min, and the supernatant was discarded. The same staining procedure described for 0 h was performed for the treated cells. Data were acquired using a BD FACSVerse flow cytometer and analyzed using FlowJo V10 software. Two biological replicates were prepared for each condition, and each sample was counted twice, with 10,000 events acquired per run.

### Toxicity assessment by analyzing hematological and biochemical parameters

C57BL/6J mice (4-6 weeks old) were intraperitoneally administered 100 µL of **1a** at a dose of 200 mg/kg or 300 mg/kg daily once for five days, or 200 mg/kg once daily for ten days. Additional groups received 100 µL of vehicle DMSO (D), **1b** or **1c** at 200 mg/kg. Mice were monitored for body weight and health regularly. After 24 h of the final dose, blood samples were collected *via* the retro-orbital plexus. A portion of the whole blood was collected in EDTA-coated tubes for complete blood cell analysis. The remaining blood was centrifuged (1000 × g, 10 min) to obtain serum, which was used for the analysis of liver enzyme and kidney protein levels using a clinical chemistry analyzer.

### RNA isolation and real-time qPCR analysis

B16F10 cells were seeded at a density of 5.0 × 10^4^ cells/mL in 10 mL of complete medium in 100 mm diameter tissue-culture treated dishes. Cells were either left untreated (U) or treated with vehicle (D), **1a**, **1b**, **1c**, or **1d** (100 μM) for 48 h. Following incubation, the media was aspirated. Cells were detached by incubating with 0.25 % trypsin (1.0 mL) and diluted with complete media (9.0 mL). The suspension was centrifuged at 500 × g for 5 min at RT, and supernatants were discarded. RNA was isolated using the TRIzol method (45). The cell pellets were resuspended in 750 μL of TRIzol reagent and incubated for five min to ensure complete dissociation of nucleoprotein complexes. Subsequently, 150 μL of chloroform was added, incubated for three min, and centrifuged at 12000 × g for 15 min at 4 °C. The colorless upper aqueous phase containing the RNA was transferred to a fresh tube. To precipitate RNA, 375 μL of isopropanol was added and mixed by pipetting. After a 10 min incubation at RT, samples were centrifuged at 12000 × g for 10 min at 4 °C, yielding a white gel-like pellet. The supernatant was discarded, and the pellet was washed by resuspension in 750 μL of 75 % aq. ethanol. Samples were vortexed briefly and centrifuged at 7,500 × g for five min at 4 °C. The supernatant was discarded, and the pellet was air-dried for 10 min in a fume hood. Finally, the RNA was resuspended in 50 μL of RNAse-free water, incubated at 55 °C for 15 min, and stored at -80 °C. RNA concentration and purity (260/280 ratio) were measured using a Biotek reader (1 μL sample volume). RNA integrity was verified by 1.0 % agarose gel electrophoresis. Subsequently, 1.0 μg of RNA was reverse transcribed into cDNA using a cDNA synthesis kit (Invitrogen, 18080051). Quantitative real-time (RT-qPCR) was performed on a QuantStudio 6 Flex system using SYBR fast and gene-specific primers (**Table 2**) to amplify melanosomal and housekeeping genes. The cycle number crossing the threshold was defined as the threshold cycle (Ct) (46). Gene expression was normalized by calculating ΔCt (Ct of target melanosomal genes – Ct of housekeeping gene). Relative quantification between GalNAc analogue-treated cells and vehicle-treated controls was performed using the comparative Ct method (2^-^ΔΔCt) (47). The fold change relative to the vehicle control (D) was plotted.

**Table 2.**
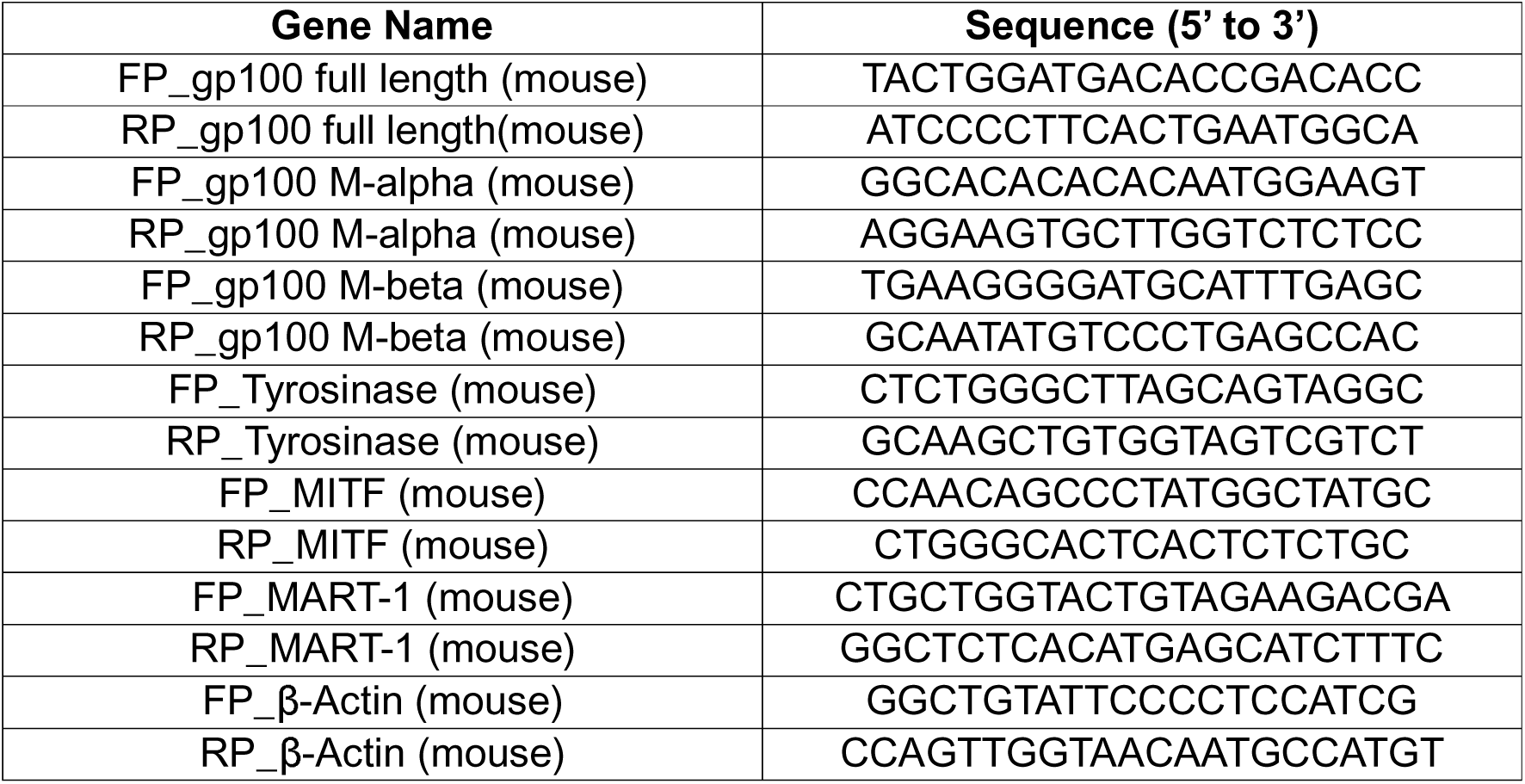
List of nucleotide sequences of primers for melanosomal-specific genes used in RT-qPCR analysis.

### RNA sequencing, library preparation, and data analysis

B16F10 cells were treated with D or **1a** (100 µM, 48 h) as described above. Following treatment, total RNA was extracted using the TRIzol method as previously described. RNA concentration and quality were assessed using a spectrophotometer. The isolated RNA was then subjected to high-throughput sequencing to obtain expression data.

The concentration of isolated RNA was quantified using a Qubit 4 Fluorometer with the Qubit HS RNA Assay kit, suitable for concentrations ranging from 0.2 to 200 ng/µL. RNA integrity was assessed using and Agilent 5200 Fragment Analyzer, and an RNA Quality Number (RQN) score was assigned to each sample. Only samples with an RQN value > 8 were selected for library preparation. RNA sequencing libraries were constructed using the QIAseq FastSelect RNA Library Kit, which features cDNA barcoding and rRNA removal, starting with 1 ng to 1000 ng of total RNA. Following construction, the quality of the final libraries was verified on the Fragment Analyzer. The libraries were then pooled and sequenced on an Illumina NextSeq 2000 platform.

Raw sequencing data was processed using fastp to perform quality control and trimming, generating high-quality trimmed data. The processed reads were then aligned to the reference Ensembl mouse genome using HISAT2. Gene expression levels were quantified by assigning mapped reads to genomic features using featureCounts. Differential expression (DE) analysis was subsequently performed using DESeq2 package to identify genes with significant expression changes between experimental conditions. Downstream analysis, including pathway enrichment and visualization, was conducted using various R packages. DEGs were identified and filtered out with the following criteria: false discover rate < 0.05 and |log2Fc| > 1.5.

### PCR Array analysis for genes involved in cancer metastasis

B16F10 cells were seeded at a density of 5.0 × 10^4^ cells/mL in 10 mL of complete medium in 100 mm diameter tissue-culture treated dishes. Cells were treated with either vehicle (D) or **1a** (100 μM) for 48 h while in suspension. Following incubation, RNA was isolated and converted to cDNA (starting with approximately 1 µg of RNA) as described previously. The prepared cDNA (50 ng per sample) was analyzed using mouse tumor metastasis RT^2^ Profiler PCR array plates. These plates contained pre-coated primers for 84 selected genes involved in metastasis, including those regulating cell adhesion, extracellular matrix components, cell cycle, cell growth and proliferation, apoptosis, and transcription factors (**Table 3, in Supplementary material**). For the assay, 50 ng/μL cDNA was added to each well, and quantitative real-time PCR was performed using the KAPA SYBR fast qPCR kit (KAPA Biosystems, KK4602). Differences between **1a**-treated cells and vehicle-treated controls were quantified using the comparative Ct method (2^-^ΔΔCt), as mentioned previously. Results were visualized as a Volcano plot, displaying the relationship between fold change and statistical significance. The x-axis represents fold change calculated as log2(2^-^ΔΔCt), and the y-axis represents statistical significance calculated as -log10(P-value)(48).

**Table 3.**
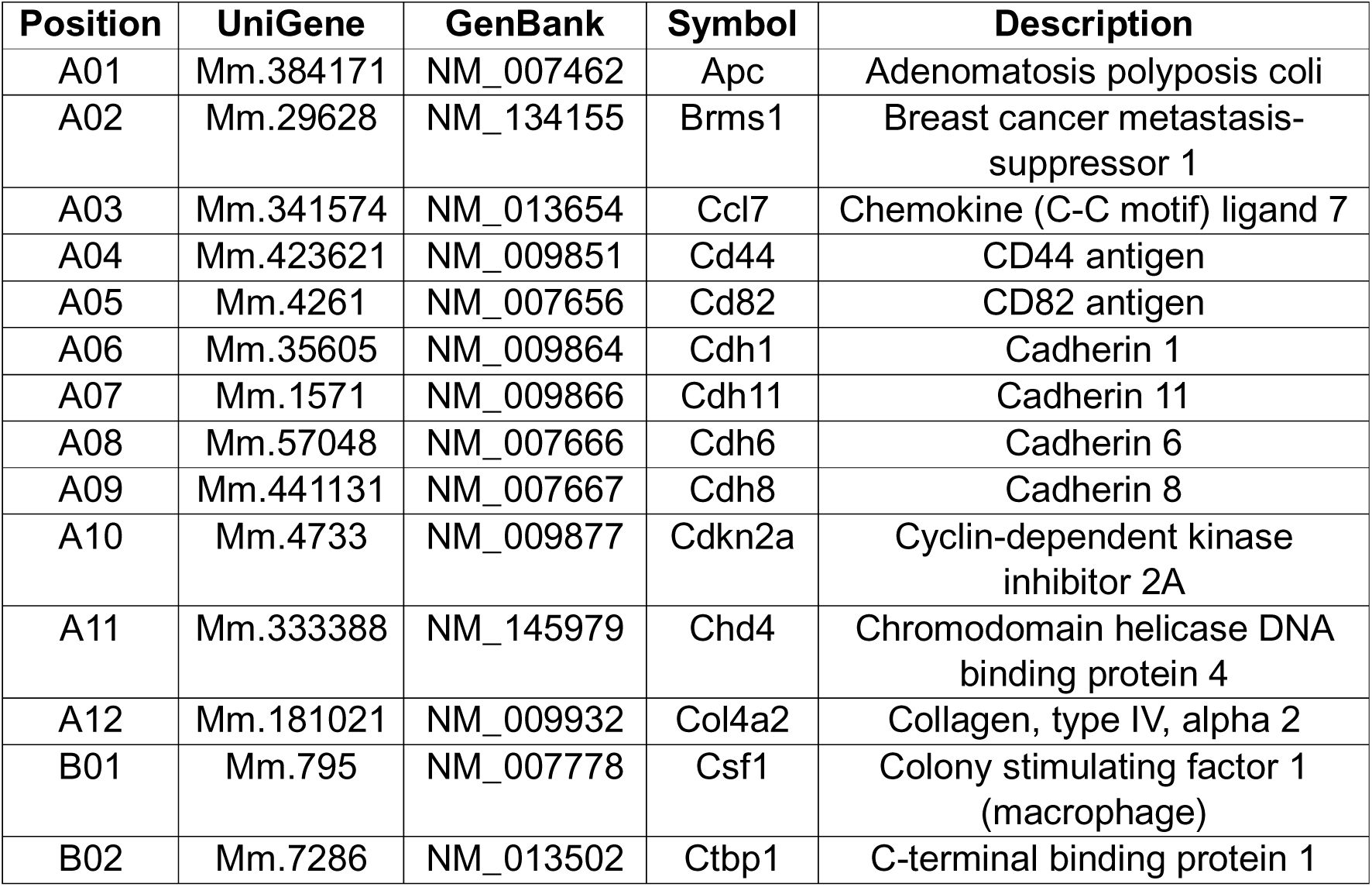
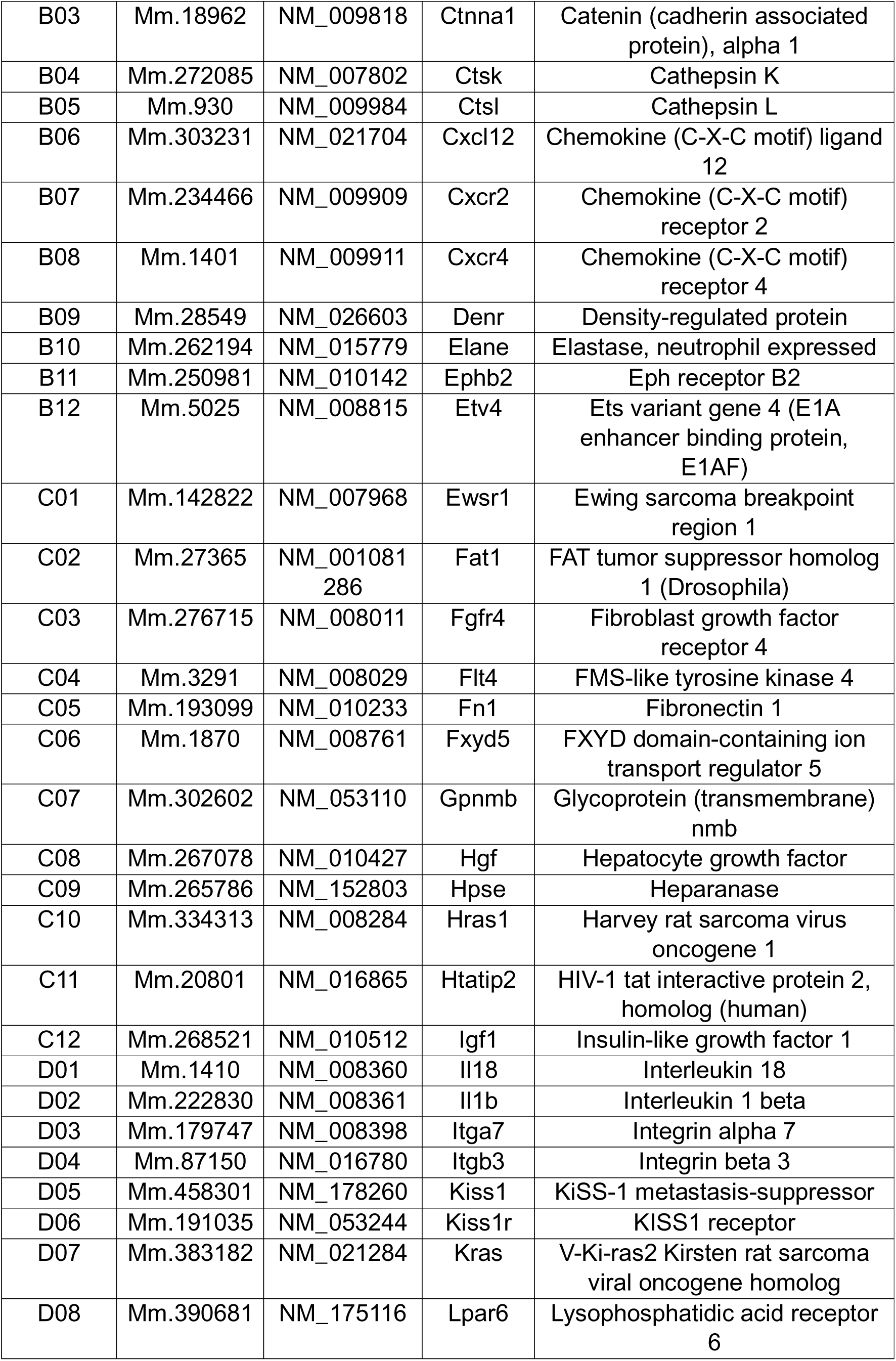
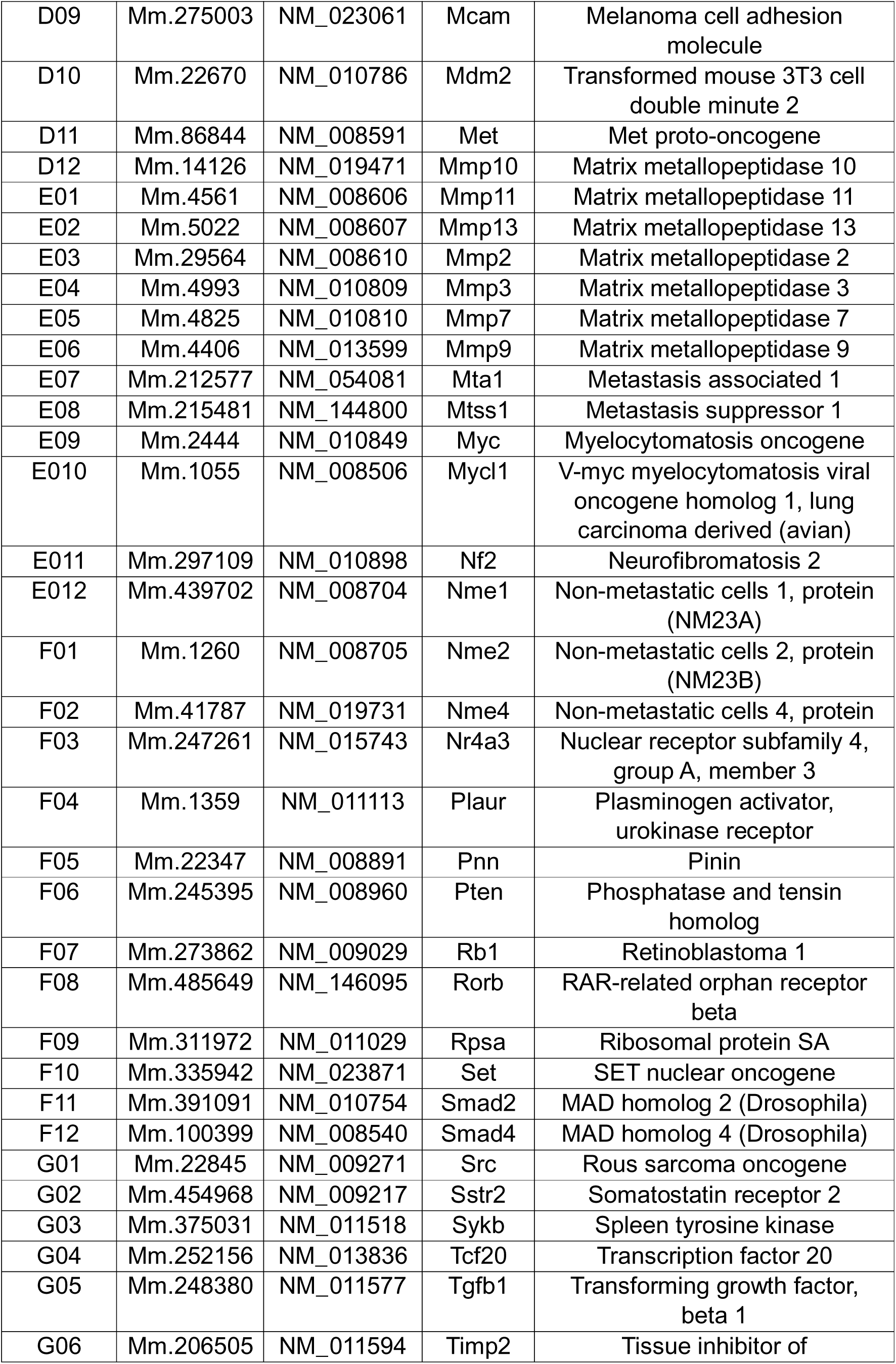
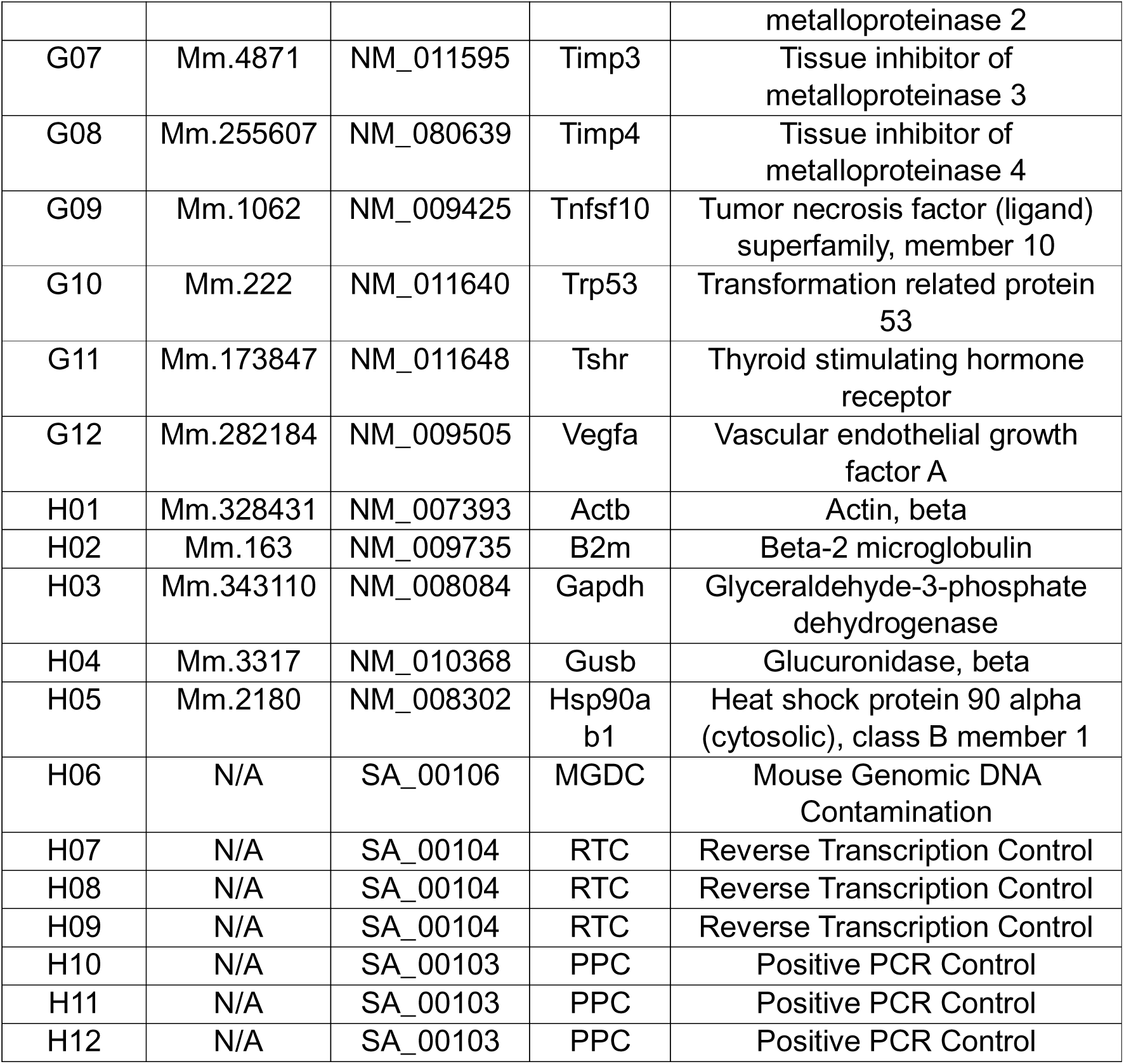
List of housekeeping (control) and metastasis-related genes included in the selected PCR array, with descriptions and GenBank IDs.

### Statistical Analysis

Statistical analyses were performed using GraphPad Prism version 8.4.3 (GraphPad Software, San Diego, USA). Data are presented as mean ± standard deviation (SD). For datasets with a Gaussian distribution, comparisons between two groups were analyzed using multiple t-tests corrected with the Holm-Sidak method. Comparisons among multiple groups were analyzed using one-way or two-way ANOVA followed by Tukey’s multiple comparisons test. For datasets with non-gaussian distribution, comparisons between multiple groups were analyzed using Kruskal-Wallis test with Dunn’s multiple comparisons test. Survival analysis was determined using the log-rank (Mantel-Cox) test. A p-value of less than 0.05 was considered statistically significant in most cases. Levels of significance are indicated in the figure legends where applicable.

## Reagents, Antibodies, and Chemicals

List of reagents used in this study:

| List of reagents | Source | Identifier (Cat. /Ref.) |
| --- | --- | --- |
| Cell lines |  |  |
| B16F10 | Dr. R. Gokhale, NII | -- |
| B16F10-Luc2 | Dr. S. Singha, NII | -- |
| Peracetyl HexNAc Analogues |  |  |
| Ac<sub>5</sub>GalNTGc (1a) | In-house |  |
| Ac<sub>4</sub>GalNAc (1b) | -do- |  |
| Ac<sub>5</sub>GalNGc (1c) | -do- |  |
| Ac<sub>3</sub>BnGalNAc (1d) | -do- |  |
| Ac<sub>4</sub>GalNAz (1e) | -do- |  |
| Ac<sub>5</sub>GlcNTGc (2a) | -do- |  |
| Ac<sub>4</sub>GlcNAc (2b) | -do- |  |

| Antibodies |  |  |
| --- | --- | --- |
| Rabbit monoclonal anti-melanoma gp100 antibody (N-term) | Abcam | ab137078 |
| Mouse monoclonal anti-melanoma gp100 antibody (HMB45) | -do- | ab787 |
| Rabbit monoclonal anti-melanoma gp100 Alexa Fluor 647 antibody (EP4863(2)) | -do- | ab246730 |
| Goat polyclonal anti-melanoma gp100 (C-term) | Origene | TA302608 |
| Rabbit monoclonal anti-tyrosinase antibody (369-377) | GenScript | RP10270 |
| Anti-β-actin IgG antibody, mouse monoclonal (AC-15) | Sigma-Aldrich | A1978 |
| Mouse IgG1 kappa isotype control | Invitrogen | 14-4714-85 |
| Goat anti-mouse IgG (H+L) Alexa Fluor 488 | -do- | A32723 |
| Goat anti-rabbit IgG (H+L) Alexa Fluor 488 | -do- | A32731 |
| Rabbit polyclonal anti-MART-1 | -do- | PA5-120982 |
| Cy5 affiniPure donkey anti-mouse IgG (H+L) | Jackson Laboratories | 715-175-150 |
| Recombinant proteins and lectins |  |  |
| FITC-avidin | Sigma | A2901 |
| Recombinant Siglec E/Fc chimera, sourced from a mouse myeloma cell line, NS0-derived mouse Siglec-E protein | R&D systems | 5806-SL |
| Biotinylated *Maackia amurensis*-II (MAL-II) | Vector Laboratories | B-1265 |
| Fluorescein isothiocyanate conjugated *Sambucus nigra* agglutinin (FITC-SNA) | -do- | FL-1301 |
| Fluorescein isothiocyanate conjugated *Vicia villosa* agglutinin (FITC-VVA) | -do- | FL-1231 |
| Fluorescein isothiocyanate conjugated wheat germ agglutinin (FITC-WGA) | -do- | FL-1021 |
| Neuraminidase from *Vibrio cholerae* | Merck (Roche) | 11080725001 |
| Chemicals and other resources |  |  |
| Doxorubicin HCl | Goldbio | D-490-500 |
| Dulbecco's modified Eagle's medium (DMEM) | Gibco | 11965-092 |
| Fetal bovine serum (FBS) | -do- | 10082-147 |
| Penicillin-Streptomycin (Pen-Strep) | -do- | 15140-122 |
| 0.25 % Trypsin-EDTA | -do- | 25200-056 |
| Blasticidin S HCl | GoldBio | B-800-100 |
| Protease inhibitor cocktail (PIC) | Sigma-Aldrich | P8340 |
| radioimmunoprecipitation assay (RIPA) buffer | -do- | R0278 |
| Propidium iodide (PI) | -do- | P4170 |
| (3-(4,5-Dimethylthiazol-2-yl)-2,5-diphenyltetrazolium bromide (MTT) | -do- | M2128 |
| DAPI | -do- | D9542 |
| Rhodamine phalloidin | Invitrogen | R415 |
| BSA | -do- | A7030 |
| Gelatin | -do- | 48722 |
| Hydrochloric acid | -do- | 30721 |
| *N*-acetyl-D-neuraminic acid | -do- | A0812 |
| Ammonium persulfate | -do- | A3678 |
| Dithiothreitol (DTT) | -do- | 43819 |
| Tris(2-carboxyethyl) phosphine hydrochloride (TCEP) | -do- | C4700 |
| β-mercaptoethanol | -do- | M7522 |
| Dimethyl sulfoxide (DMSO) | Fisher Scientific | 67-68-5 |
| Melanin | Merck | M0418 |
| Sodium hydroxide | -do- | 1.93102.0521 |
| Sodium chloride | -do- | 1.93206.0521 |
| Sodium thiosulfate pentahydrate | -do- | 61778405001730 |
| Sodium carbonate anhydrous | Qualikems | S039112 |
| Di-sodium hydrogen phosphate dihydrate | Merck | 1.93622.0521 |
| Potassium dihydrogen phosphate | -do- | 7778-77-0 |
| Potassium chloride | -do- | 7447-40-7 |
| Tris(hydroxymethyl)aminomethane | -do- | 1.93315.0521 |
| Glycerol anhydrous | -do- | 1.07051.0521 |
| Glycine | -do- | 1.94907.0521 |
| Tween 20, | -do- | 8.22184.0521 |
| Paraformaldehyde powder | -do- | 158127 |
| Chloroform | -do- | 60244410001730 |
| 2-propanol | -do- | 1.94524.0521 |
| Methanol | -do- | 60600725001730 |
| Glacial acetic acid | -do- | 1.93402.0521 |
| 37 % aq. Formaldehyde | -do- | 1.94989.0521 |
| Ethanol absolute | -do- | 64-17-5 |
| Crystal violet | -do- | C0775 |
| RT<sup>2</sup> Profiler PCR Array Mouse Tumor Metastasis kit | Qiagen | PAMM-028ZA-6 |
| Nuclease-free water | -do- | 1039498 |
| Carboxyfluorescein succinimidyl ester (CFSE) | Invitrogen | C1157 |
| TRIzol reagent | -do- | 15596-026 |
| SuperScript III First-Strand Synthesis System | -do- | 18080051 |
| eFluor 506 fixable viability dye | -do- | 65-0866-14 |
| ProLong Gold Anti-fade | Life Technologies | P36934 |
| Skim milk powder | HiMedia | GRM1254 |
| Bradford reagent | Bio-Rad | 5000006 |
| 40 % aq. Acrylamide solution | -do- | 1610140 |
| 2 % aq. Bis acrylamide solution | -do- | 1610142 |
| Precision Plus Protein Dual Color Standards | -do- | 1610374 |
| Bromophenol blue electrophoresis purity reagent | -do- | 161-0404 |
| Triton X-100 Detergent | -do- | 1610407 |
| Ethidium Bromide solution | -do- | 161-0433 |
| Sodium dodecyl sulfate (SDS) | SRL Chem | 14374 |
| Anhydrous sodium carbonate | -do- | 1949157 |
| *N, N, N', N'* Tetramethyl-ethylenediamine (TEMED) | MP Biochemicals | 194019 |
| Agarose Plus | Pronastar | 8045 |
| 1kb DNA ladder | New England Biolabs | N3232S |
| 6× Gel Loading Dye (blue) | -do- | B7021S |
| KAPA SYBR FAST | KAPA biosystems | KK4602 |
| DBCO-Cy5 | Click Chemistry Tools | A130-25 |
| Maleimide-PEG<sub>3</sub>-biotin | -do- | 1029 |
| Silver nitrate | BDH | I132529 |
| D-Luciferin potassium salt | GoldBio | LUCK |
| Ketamine (80 mg/kg) | Zokent, Miracalus Pharma Private Limited | -- |
| Xylazine (20 mg/kg) | Indian Immunologicals Limited | -- |
| 0.2 µm syringe PTFE filters | Millex-LG | SLLG013SL |
| PET membrane inserts 24 well 8.0 µm pore size | Corning | 353097 |
| Matrigel coated insert 24 well 8.0 µm pore size | -do- | 354480 |
| 1 mL Syringe (26G ½) | BD | 301321 |
| Digital vernier calipers | SKADIOO | -- |

## Supplemental Material

Supplementary material includes supporting results, discussion, methods (in the main text and methods), figures S1−S5, and Table 3 with corresponding legends (in the supplementary section).

## Data availability

The data supporting the findings of this study are available from the corresponding author upon reasonable request.

## Acknowledgements

We thank Mr. Aryan, Ms. Riddhima, Mrs. Jyoti, Dr. Shubham, and Mrs. Monica for providing the HexNAc analogues used in this study. We also thank Mr. Anand and Mr. Aslam for providing autoclaved PBS, tips, plasticware, glassware, and all the technical help. We are grateful to Mrs. Archana, Mr. Arun, and Mr. Shahnawaz at the Central Instrumentation Facility (CIF) for their technical assistance. We acknowledge Mr. Athar and Dr. Devram from IIL, NII, for their support in accessing and using the Nikon inverted microscope. We sincerely acknowledge Sureshkumar Venkadesan from the Bioinformatics Centre, NII, for his valuable assistance with NGS RNA sequencing data analysis. We also thank the NGS Sequencing Facility at NII for their support with RNA sequencing services.

## Author contributions

TJ, AR, SGS, and AiR conceptualized the study, designed the experiments, analyzed the data, and wrote the manuscript. TJ, AR, SGK, and AiR performed the experiments. AR performed experiments *in vivo* in mice. SS provided critical inputs and the cell lines. NP trained and supervised the experiments in mice. SGS directed and supervised the research. All authors reviewed and approved the final manuscript.

## Supplementary Material

**Figure S1.**
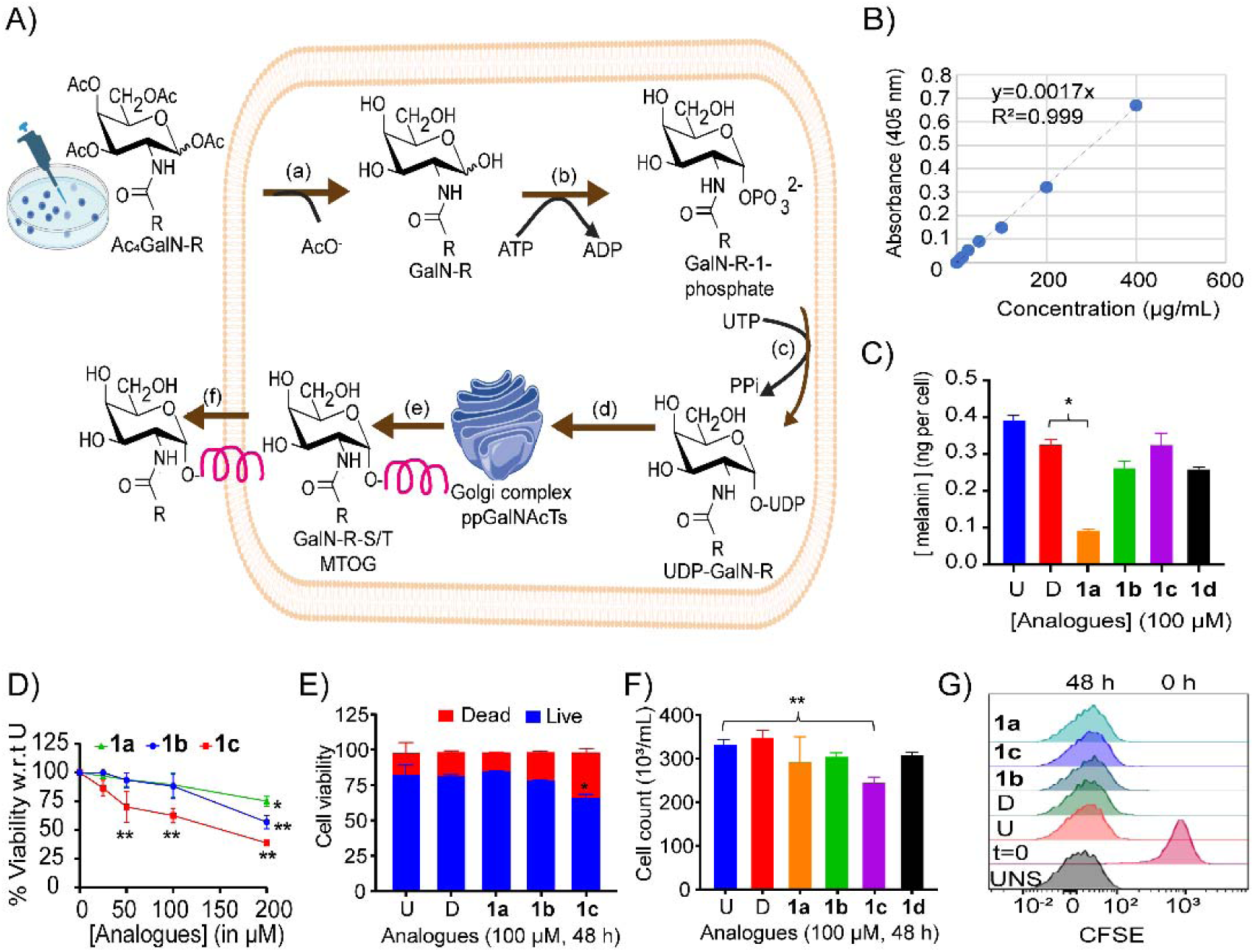
Metabolic glycan engineering (MGE) by GalNAc analogues selectively suppresses melanogenesis without affecting cell viability or proliferation. (A) Schematic illustrating MGE by peracetylated GalNAc analogues. (a) Peracetylated analogues passively diffuse into cells and are deacetylated by intracellular esterases. (b) The resulting free analogue undergoes phosphorylation at the C1-OH position by GalNAc-1-kinase. (c) Phosphorylated intermediates are activated to UDP-GalNAc analogues by UDP-GalNAc pyrophosphorylase. (d) These donors are transported into the Golgi lumen via the UDP-GalNAc transporter and are subsequently utilized by the mucin-type *O*-glycosylation enzymatic machinery. (e) This results in the incorporation of modified GalNAc analogues into *O*-glycans. (f) The modified *O-*glycans are expressed on cell surface glycoproteins, enabling detection and functional analyses. (B-C) Effect of GalNAc analogues on melanin biosynthesis. (B) Melanin standard curve generated using synthetic melanin solutions ranging from 0 to 400 µg/mL. (C) Quantification of intracellular melanin content in B16F10 cells upon analogue treatment. Data is presented as melanin concentration (ng per cell). Error bars represent SD of three BR. Statistical significance was determined using one-way ANOVA. P-values are *<0.05 compared with vehicle D. (D-G) Effect of GalNAc analogues on B16F10 cell viability and proliferation. (D) Cell viability assessed by MTT assay upon analogue treatment (0-200 µM), expressed as percentage relative to untreated cells. Seeding density: 10,000 cells/well (E) Percentage of dead and live cells determined by trypan blue exclusion following analogue treatment. Approximately 100 cells per sample were counted using a hemocytometer. (F) Cell density (× 10^3^/mL) measured after 48 h upon analogue treatment at 100 µM. Cells were plated at 5.0 × 10^4^/mL in 10 mL of DMEM. (G) CFSE dye dilution assay assessing cell proliferation. Representative histograms show CFSE fluorescence at 0 h and dilution at 48 h upon analogue treatment (100 µM, 48 h) in B16F10 cells. Error bars represent SD of three BR in (D-F) and two BR (each sample counted twice for 10,000 PI-negative events) in (G). Statistical analysis was performed using two-way ANOVA for MTT, Trypan blue, and CFSE assays, and one-way ANOVA for cell count. P-values are *<0.05, **<0.01, compared to vehicle. U, untreated; D, DMSO (vehicle); **1a**, Ac_5_GalNTGc; **1b**, Ac_4_GalNAc; **1c**, Ac_5_GalNGc; and **1d**, Ac_3_BnGalNAc.

**Figure S2.**
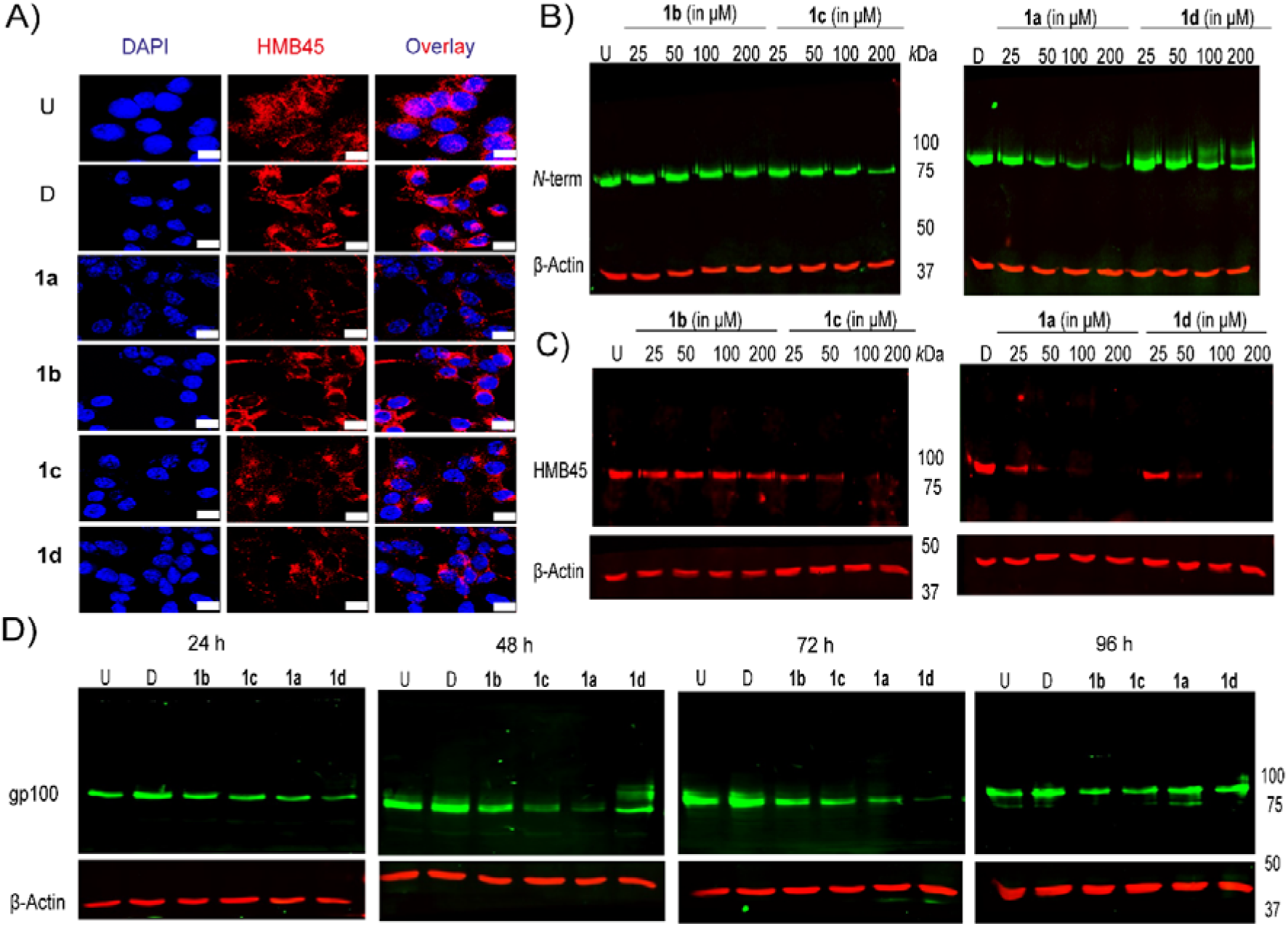
Dose-dependent and time-course modulation of PMEL/gp100 glycosylation by GalNAc analogues. (A) Confocal imaging of B16F10 cells treated with GalNAc analogues (100 µM, 48 h), fixed, permeabilized, and stained with anti-gp100 antibody (HMB45, red), along with DAPI (blue). Imaging was performed using 63× oil immersion on a Zeiss LSM 980 confocal microscope. Images shown are representative of three BR with two technical replicates per BR. Each replicate was conducted on one cover slip with five fields imaged per coverslip. Scale bar: 5 µm. (B-C) B16F10 cells were treated with GalNAc analogues (0-200 µM, 48 h). Cells were lysed, and cell lysates were subjected to western blotting using: (B) anti-gp100 (*N*-term) antibody; (C) anti-gp100 (HMB45) antibody, which specifically recognizes a glycan-dependent epitope on PMEL/gp100. The blots showed dose-dependent loss of HMB45 reactivity upon **1a** treatment. (D) Time-course analysis of PMEL/gp100 modulation. B16F10 cells were treated with GalNAc analogues (100 µM) and lysed at respective time points (24, 48, 72, and 96 h). Cell lysates were probed with anti-gp100 (*N*-term) antibody. Western blots showed maximal inhibition of PMEL/gp100 levels at 48 h upon **1a** treatment, with recovery of PMEL/gp100 levels by 96 h, consistent with the reversible nature of MGE. β-actin blots are shown as loading controls. Blots shown are representative of two BR with two technical replicates per BR. U, untreated; D, DMSO (vehicle); **1a**, Ac_5_GalNTGc; **1b**, Ac_4_GalNAc; **1c**, Ac_5_GalNGc; and **1d**, Ac_3_BnGalNAc.

**Figure S3.**
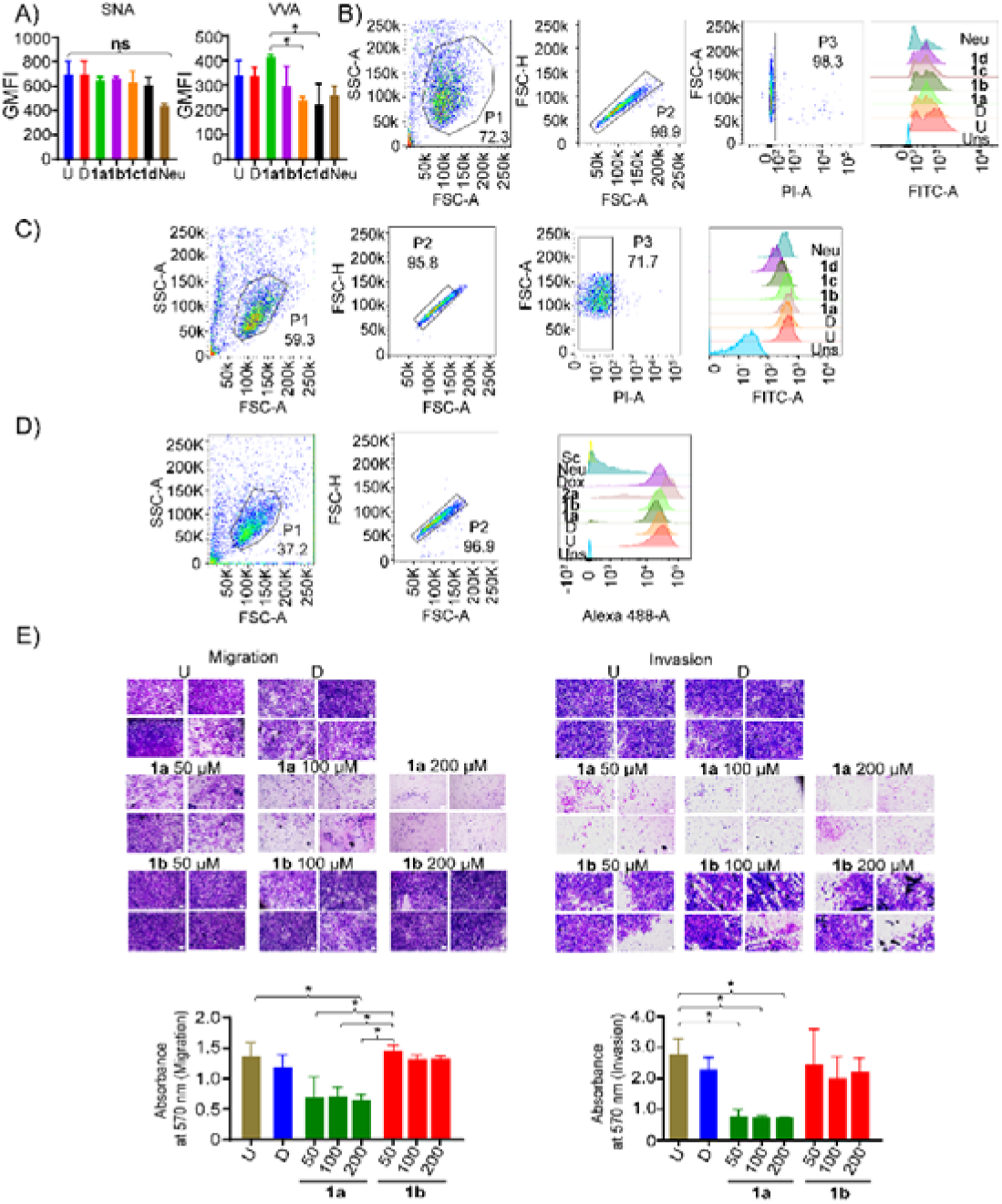
1a-mediated decrease in cell-surface sialylation, Siglec ligand engagement, and metastatic traits in melanoma cells. (A-C) Quantification of cell-surface sialic acid levels in B16F10 cells treated with GalNAc analogues. (A) B16F10 cells treated with GalNAc analogues (100 µM, 48 h) were stained with fluorescein-conjugated SNA (0.5 µg/mL) and VVA lectin (10 µg/mL) for 15 min at RT. Cells treated with Neuraminidase from *Vibrio cholerae* (20 milliunits/mL) for 30 min were used as negative controls. Stained cells were analyzed by flow cytometry, and GMFI of FITC-positive populations was quantified and plotted as bar graphs. Error bars indicate the SD of two BR with two technical replicates each. Statistical significance was determined using Kruskal-Wallis test followed by Dunn’s multiple comparisons test. P-value * ≤0.05, and all other comparisons were non-significant. (B-C) Flow cytometry gating strategy for lectin binding analysis. Cells were sequentially gated on SSC-A vs FSC-A (to exclude debris), FSC-H vs FSC-A (to select single cells), and FSC-A vs PI-A (to select live cells). Histograms represent FITC-positive populations for (B) SNA binding and (C) VVA binding. (D) Gating strategy for Siglec-E-Fc binding. Cells were sequentially gated on SSC-A vs FSC-A, and FSC-H vs FSC-A to select single cells. Histograms depict AF488-positive populations corresponding to Siglec-E-Fc binding. (E) **1a**-treatment suppresses melanoma cell migration and invasion in a dose-dependent manner. B16F10-Luc2 (1.0 10^5^) cells were treated with vehicle (DMSO), **1a**, or **1b** at 50, 100, or 200 µM concentrations and seeded into transwell inserts. For invasion assays, inserts were coated with Matrigel to mimic the extracellular matrix. After monolayer formation (∼12 h), a serum gradient was introduced, and cell migration or invasion through an 8.0 μm PET membrane was assessed after 48 h. Representative images of cells that migrated or invaded to the bottom side of the membrane, fixed and stained with 0.25 % crystal violet, are shown. [Scale bar: 100 µm, images representative of four fields per insert, 10× objective]. Cells on the lower surface were solubilized in DMSO, and absorbance was measured at 570 nm. Error bars represent the SD of two BR. Statistical significance was determined using one-way ANOVA with Tukey’s multiple comparisons test. P-value * ≤ 0.1 for invasion and P-value * ≤ 0.05 for migration. U, untreated; D, DMSO (vehicle); **1a**, Ac_5_GalNTGc; **1b**, Ac_4_GalNAc; **1c**, Ac_5_GalNGc; **1d**, Ac_3_BnGalNAc; **Neu**, neuraminidase; **2a**, Ac_5_GlcNTGc; **Dox**, Doxorubicin; IC, isotype control; and Uns, Unstained.

**Figure S4.**
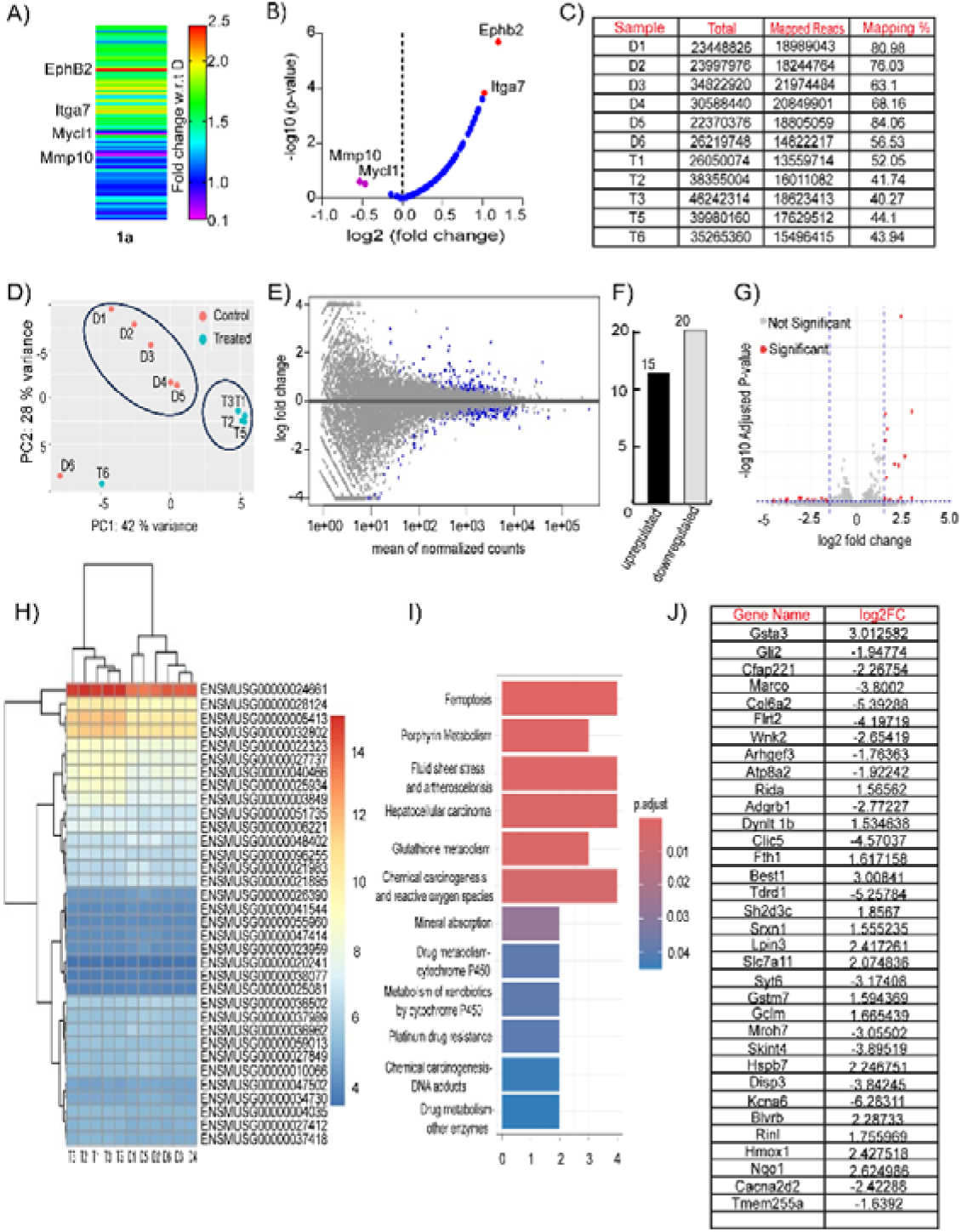
Targeted metastasis gene profiling reveals selective transcriptional effects of 1a, while global transcriptomic analysis indicates minimal off-target gene regulation. (A-B) Changes in mRNA levels of metastasis-associated genes in B16F10 cells following **1a** treatment assessed using a mouse metastasis RT^2^ Profiler PCR array. (A) Heatmap showing fold changes in metastatic genes expression upon **1a** treatment (100 µM, 48 h) relative to the vehicle (DMSO). Four significantly altered genes are highlighted: EphB2 and ITGA7 were upregulated by 2-fold, while MMP-10 and Mycl-1 were downregulated by 0.5-fold upon **1a** treatment. (B) Volcano plot shows relative expression changes, depicted as log2(fold change), plotted against statistical significance -log10(P-value). Red indicates genes significantly upregulated by ≥2-fold. Pink indicates genes significantly downregulated by ≥2-fold upon 1a treatment. Data represent n = 2 BR run on two independent PCR array plates. (C-J) Transcriptomic profiling of B16F10 cells treated with **1a** reveals minimal global gene expression changes. (C) Summary of sequencing read statistics, including total reads, mapped reads, and mapping efficiency for all qualified samples. Sample T4 was excluded due to low RNA quality (Status: Fail). (D) Principal Component Analysis (PCA) plot showing distinct clustering between the control (DMSO, red) and treated (**1a**, blue) groups. (E) MA plot visualizing log2 fold changes versus mean normalized expression levels. (F) Bar graph summarizing differentially expressed genes (DEGs), with 15 upregulated and 20 downregulated genes. (G) Volcano plot displaying DEGs; red dots represent genes meeting the significance threshold (adjusted P-value < 0.05 and |log2FC| > 1.5), while grey dots represent non-significant genes. (H) Heatmap with hierarchical clustering of significant DEGs across all samples. (I) Pathway enrichment analysis (KEGG/GO) of DEGs, highlighting enrichment of pathways related to ferroptosis, porphyrin metabolism, and glutathione metabolism. Bars are colored based on adjusted P-value. (J) Table of DEGs listing gene name and log2 fold change (log2FC). D, DMSO (vehicle) and 1a, Ac_5_GalNTGc.

**Figure S5.**
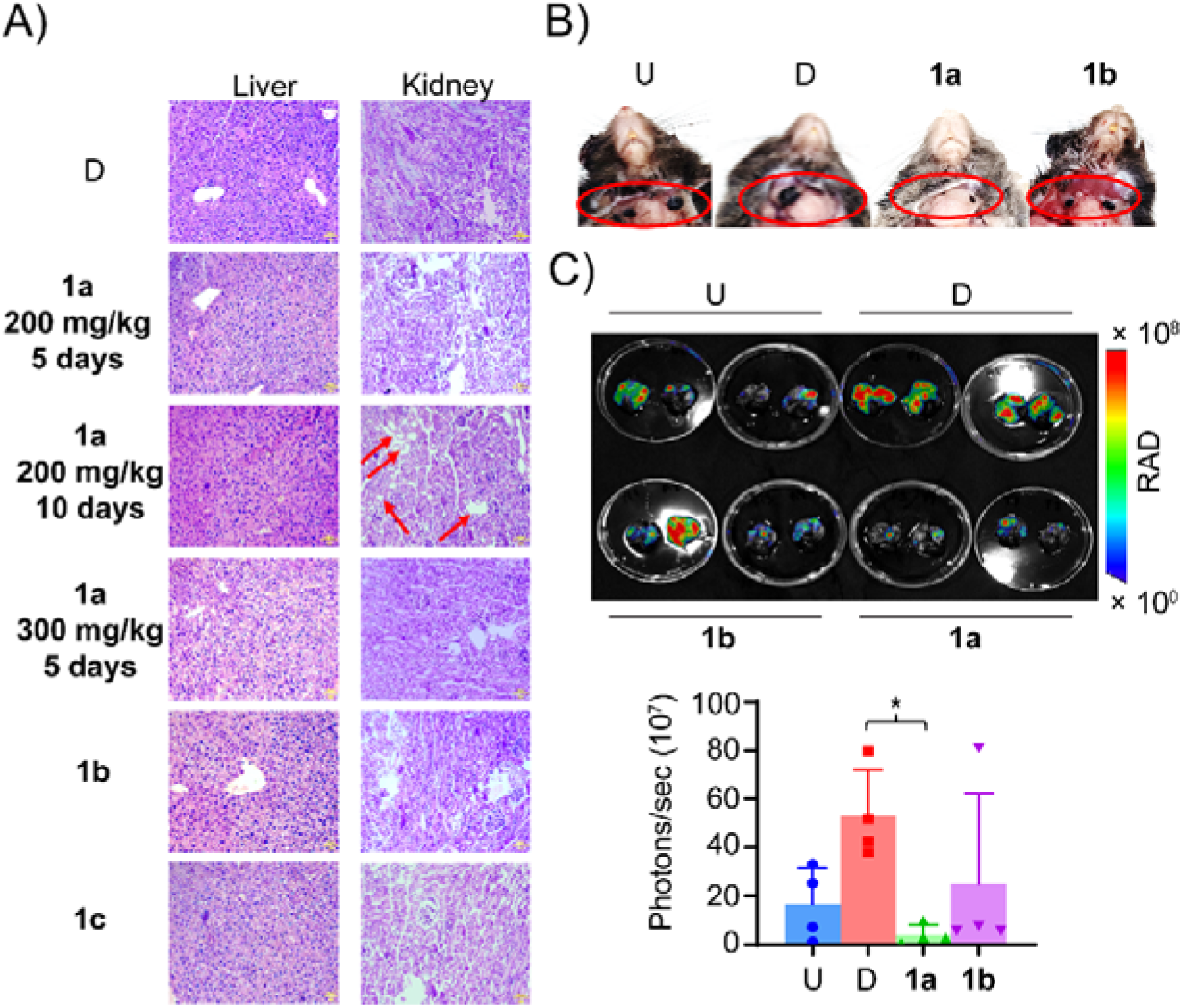
Histopathological evaluation of liver and kidney sections reveals negligible toxicity of 1a, while intravenous injection of 1a-pretreated melanoma cells slows down metastases. (A) H&E staining on mouse liver and kidney sections was analyzed after treatment with vehicle DMSO (D), **1a** at 200 mg/kg for five or ten days, 1a at 300 mg/kg for five days, **1b** or **1c** at 200 mg/kg for five days. Representative images of H&E from one mouse is shown from each group. n = four mice per group. Scale bar: 50 µm, 20× objective. (B) Metastatic nodules were also observed in the lymph nodes surrounding the neck region following dissection. (C) *Ex vivo* bioluminescence imaging of lungs soaked in D-luciferin substrate was performed using a Lago X imager. BLI (photons/sec) was quantified using Aura software. Error bars represent the SD of n = four mice per group. Statistical significance was determined using the Kruskal-Wallis test followed by Dunn’s multiple comparisons test. P-value * ≤ 0.05. U, untreated; D, DMSO (vehicle); **1a**, Ac_5_GalNTGc; **1b**, Ac_4_GalNAc; and **1c,** Ac_5_GalNGc.

